# Computational markers of rhythmic timing in Parkinson’s disease

**DOI:** 10.64898/2026.10.22.760665

**Authors:** Victor Pando-Naude, Viktor Hartmann Kuang, Karen Lambæk Christensen, Jan Stupacher, Peter Vuust, Peter E. Keller

**Affiliations:** Center for Music in the Brain, Department of Clinical Medicine, Aarhus University & The Royal Academy of Music, Aarhus/Aalborg, Denmark; Department of Neurosurgery, Aarhus University Hospital, Aarhus, Denmark; Department of Physiotherapy, VIA University College, Aarhus, Denmark; MARCS Institute for Brain, Behaviour and Development, Western Sydney University, Sydney, Australia

**Keywords:** Parkinson’s disease, rhythmic auditory stimulation, sensorimotor synchronization, timing, timekeeper noise, computational neuroscience

## Abstract

Sensorimotor synchronization requires the nervous system to predict upcoming events while continuously correcting the timing of ongoing actions. Parkinson’s disease (PD) is associated with impaired temporal processing, but it remains unclear which cognitive-computational mechanisms underlie rhythmic timing deficits. Here, we applied the Adaptation and Anticipation Model (ADAM) to synchronized finger-tapping data from participants with PD and controls across four steady tempi. After data screening, analyses included 35 participants with PD and 28 controls. PD was associated with reduced synchronization accuracy and stability, reflected in larger absolute asynchronies and higher variability of asynchronies. ADAM parameter estimates indicated lower phase correction (coupling strength) and higher central timekeeper noise in the PD group, whereas period correction (tempo matching) and peripheral motor noise showed no robust group differences. Across participants, timekeeper noise predicted synchronization accuracy. Phase correction, timekeeper noise, and motor noise predicted synchronization stability, but their associations differed between groups: phase correction was more closely associated with stability in PD, whereas timekeeper and motor noise were more closely associated with stability in controls. These findings suggest that sensorimotor synchronization deficits in PD reflect reduced use of recent timing errors to guide upcoming movements and less precise internal timekeeping. Error correction and timekeeper noise thus emerge as candidate computational markers of altered rhythmic timing in PD, with potential relevance for understanding individual differences in responses to rhythmic auditory cueing.

## 1. Introduction

Human movement is temporally organized and includes a range of rhythmic behaviours: walking, speaking, playing music, and interacting with others all require the nervous system to anticipate when events will occur and to adjust ongoing actions when prediction and sensory evidence diverge. Sensorimotor synchronization offers a particularly tractable way to study these mechanisms because the timing of external events can be precisely controlled, the timing of actions can be measured with millisecond resolution, and the resulting behaviour can be formalized computationally. Accurate movements depend on an internal timekeeper (e.g., a neural oscillator) to guide movement and at least two complementary processes: prediction of upcoming pacing events and error correction based on previous asynchronies [1]. Even when the pacing sequence is perfectly regular, synchronization is not passive entrainment, it requires the continuous use of prior timing information to maintain alignment with the external rhythm while compensating for internal variability in planning and execution [2–4].

Parkinson’s disease (PD) is classically defined by motor signs, but it also affects temporal control, action initiation, and the generation of internally timed movements [5]. This creates an apparent paradox: although PD compromises timing and movement control, external rhythmic auditory cues can improve gait in many individuals [6–9]. Rhythmic auditory stimulation (RAS) refers to the use of regular external sounds, such as metronome beats or rhythmically structured music, to support movement timing [7, 10]. Its effects suggest that RAS does not simply replace a defective motor command with an external metronome. Rather, its benefits may depend on residual timing mechanisms that allow an individual to use auditory structure for movement planning, temporal prediction, and error correction [11]. Consistent with this view, individual differences in rhythmic skills have been linked to the magnitude of gait improvement during RAS [12, 13]. Identifying the mechanisms that support these individual differences is therefore relevant both for basic models of timing and for the development of personalized rhythm-based interventions.

The Adaptation and Anticipation Model (ADAM) provides a viable computational framework for decomposing synchronization behaviour into interpretable parameters that represent different underlying mechanisms [14] (**Figure 1**). The model was developed within the broader ensemble-skills framework, where coordinated action depends on the interaction between internal models for self-generated actions and predicted external events [15–18]. In its full form, ADAM includes modules for adaptation, anticipation, and joint ‘self-other’ integration. The adaptation module implements phase correction and period correction. Phase correction refers to rapid compensation for an immediately preceding synchronization error, whereas period correction reflects adjustment of the internal period or tempo estimate if a mismatch is detected [19]. The anticipation module predicts upcoming event timing, and the joint module serves to reduce discrepancies between self-related action plans and externally predicted events before movement execution [20, 21].

**Figure 1.**
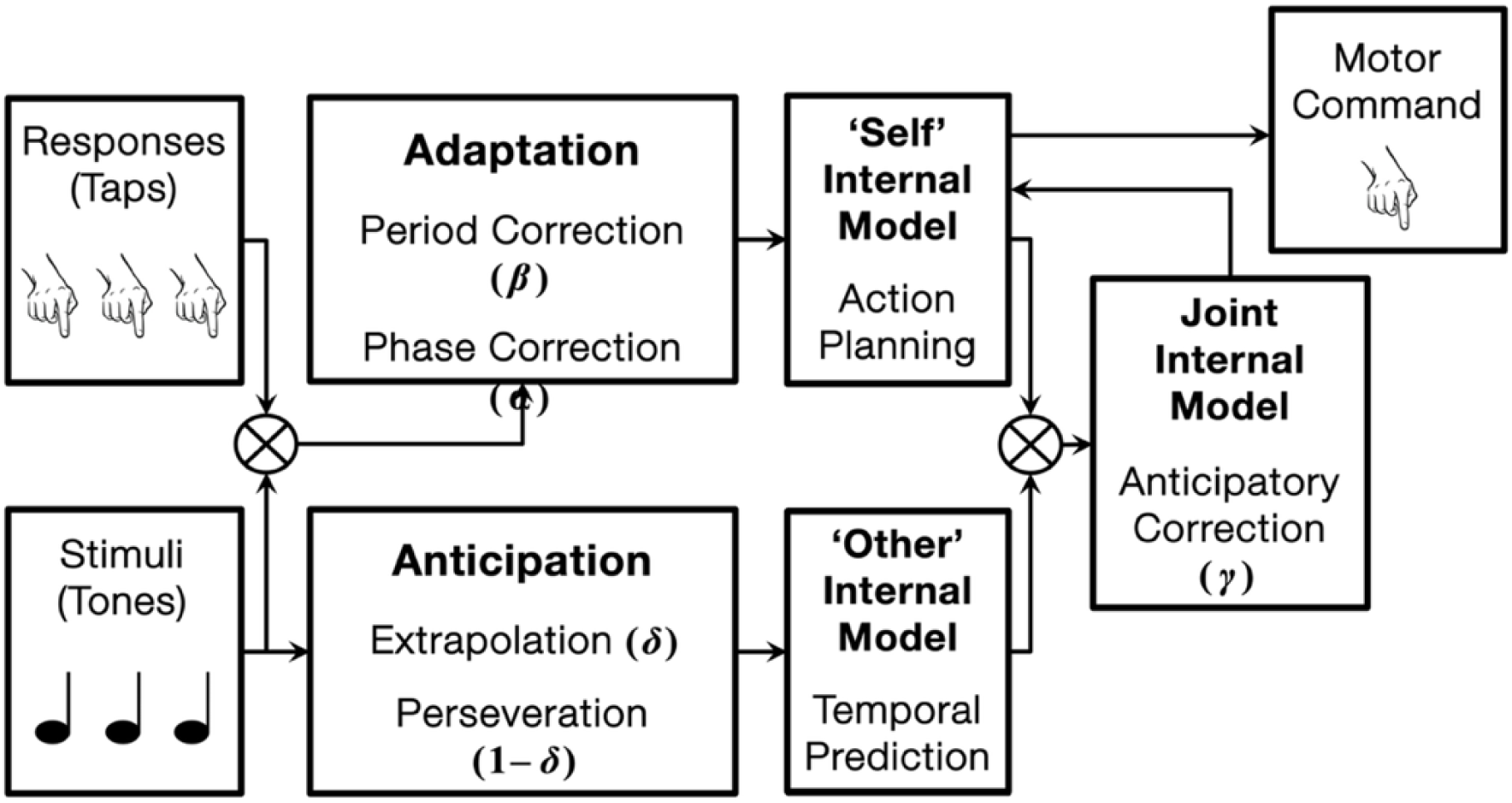
Schematic diagram of the Adaptation and Anticipation Model (ADAM) of an individual producing finger tap responses in synchrony with stimulus beats in a pacing sequence. Phase correction (α) and period correction (β) mechanisms in the ‘adaptation’ module influence action planning in a ‘self’ internal model. The ‘anticipation’ module controls the weighting (δ) of two processes, one entailing linear extrapolation of previous inter-onset intervals in the external sequence and the other copying the previous interval, which inform temporal predictions in an ‘other’ internal model. An anticipatory error correction mechanism (γ) a ‘joint’ internal model reduces discrepancies between (self) plans and (other) predictions before motor commands are issued. The processes represented in the diagram are assumed to act on internal timekeepers (which have inherent variability) and ultimately influence the timing of motor commands (which have an independent source of inherent variability). Figure adapted from [20].

A strength of ADAM is that it separates correction parameters from other variability parameters. This distinction is theoretically important in PD. Poor synchronization could arise because participants correct errors less efficiently, because their internal timekeeper is noisier, because motor execution is more variable, or because these mechanisms interact. Traditional behavioural outcomes, such as mean asynchrony or tapping variability, alone cannot distinguish these possibilities. ADAM estimates phase correction, period correction, central timekeeper noise, and motor noise from the structure of tapping time series, thereby linking observed performance to candidate computational processes [22–24] (**Figure 2**). Prior modelling work with neurological lesions suggests that basal ganglia and cerebellar disorders can affect different aspects of temporal adaptation and anticipation [24], making this approach especially relevant for PD.

**Figure 2.**
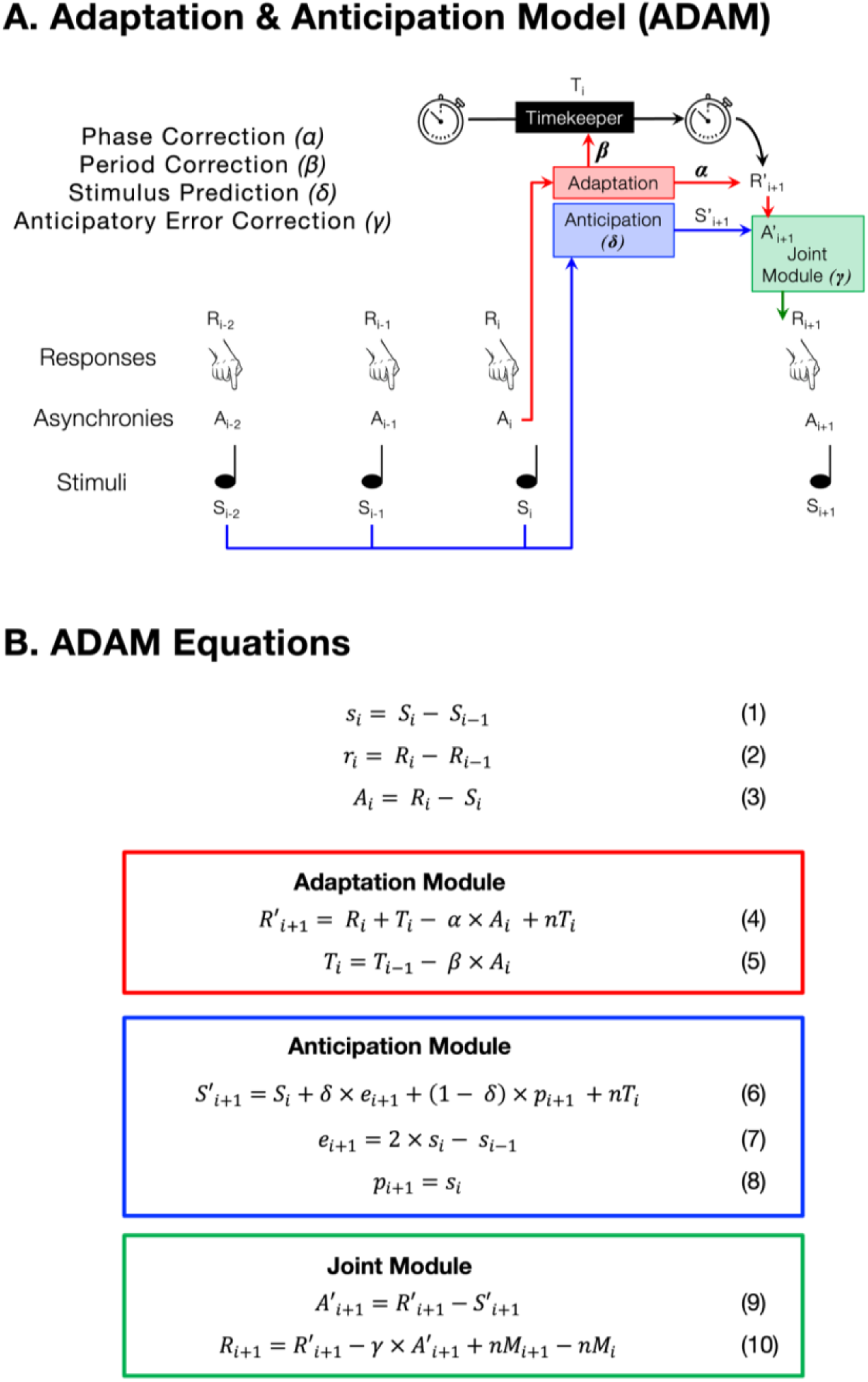
Formal components of the ADAM framework. **A)** Process diagram showing the relationship between stimuli (*S*), responses (*R*), and asynchronies between stimuli and responses (*A*). Tap response timing (e.g., *R_i+1_*) is computed based on previous stimuli, responses, an internal timekeeper, and compensatory mechanisms within the adaptation, anticipation, and joint modules. Phase correction (α) and period correction (β) in the adaptation module generate action plans for timing the next response (*R*′*_i+1_*). The weighting (δ) of extrapolation and perseveration processes in the anticipation module generate temporal predictions about the timing of the next sequence beat (*S*′*_i+1_*). Anticipatory error correction (γ) in the joint module reduces discrepancies between plans and predictions before a motor command is issued and the response is executed (*R_i+1_*). **B)** Equations for computing inter-stimulus (beat onset) intervals (1), inter-response (tap) intervals (2), and asynchronies (3) from behavioural data, which serve as input to equations for ADAM’s adaptation module (4 & 5), anticipation module (6-8), and joint module (9 & 10) used in parameter estimation. *S*, stimulus (beat) onset; *R*, response (tap) onset; *s*, inter-stimulus interval, *r*, inter-response interval, *A*, asynchrony, *T*, timekeeper interval; *R*′, planned response onset; *S*′ predicted stimulus onset; *A*′, anticipated error; *e*, extrapolated stimulus interval; *p*, perseverated stimulus interval; *nT*, timekeeper noise; *nM*, motor noise; *α*, phase correction, *β*, period correction; *δ*, prediction/tracking weight; *γ*, anticipatory error correction. Figure adapted from [20].

The present study applied ADAM to synchronized finger tapping in participants with PD and age-matched controls. Participants tapped with auditory pacing sequences at four steady tempi: 100, 110, 120, and 130 beats per minute, corresponding to inter-beat intervals of 600, ∼545, 500, and ∼462 milliseconds (ms). These tempi span a range around commonly comfortable synchronization rates [1] and allow us to ask whether group differences are constant across tempo or amplified at faster or slower pacing rates. This is clinically relevant because cueing rate is a central parameter in RAS, yet previous work suggests that the effects of auditory cues in PD are frequency-, task-, and context-dependent rather than uniformly better at slower rates [25]. However, this assumption is not straightforward: slower pacing also requires participants to maintain longer internal intervals, which may increase reliance on internal timing precision [26]. Testing a range of steady tempi therefore allowed us to ask whether PD-related synchronization deficits are specifically accentuated at slower rates or instead reflect a more general alteration of adaptive timing mechanisms. Because the sequences were steady rather than tempo-changing, the adaptation-only version of ADAM was used [20, 27]. This version provides estimates of phase correction, period correction, timekeeper noise, and motor noise [21, 22].

We expected participants with PD to show reduced synchronization performance, expressed as larger absolute asynchrony and greater variability of asynchronies. At the level of underlying mechanisms, we expected these behavioural impairments to be associated with lower period correction and increased timekeeper noise. We further asked whether tempo modulated these group differences and whether ADAM parameters explained synchronization performance similarly in participants with PD and controls. The central aim was therefore not only to describe tapping differences between groups, but to identify which components of error correction and internal timing processes best account for impaired rhythmic synchronization in PD.

## 2. Methods

Applying the ADAM to sensorimotor synchronization data from n = 35 participants with Parkinson’s Disease (PD) and n = 28 controls proceeded in three stages: (1) data pre-processing, (2) computing measures of synchronization performance and ADAM parameter estimates, and (3) data analysis.

### 2.1. Participants

The study was conducted at the Centre for Music in the Brain and the Section of Sports Science, Aarhus University, Aarhus, Denmark in collaboration with VIA University College, Aarhus, Denmark. A PD group (n = 35, age range =50-82, mean = 70.1, SD = 7.4, 13 female, 22 male) and an age-matched control group (n = 28, age range = 45–83, mean = 68.5, SD = 8.2, 18 female, 10 male) participated in this study (**Table 1**). The target sample size was 30 participants per group, informed by preliminary PD–control differences in phase correction and timekeeper noise observed in a pilot study (Cohen’s d = 0.86–0.89). Power calculations based on a two-sided independent-samples t-test with α = .05 indicated that 30 participants per group would provide 80% power to detect d = 0.74. With 24 analysable participants per group, the detectable effect at the same significance level and power would be approximately d = 0.83.

**Table 1.**
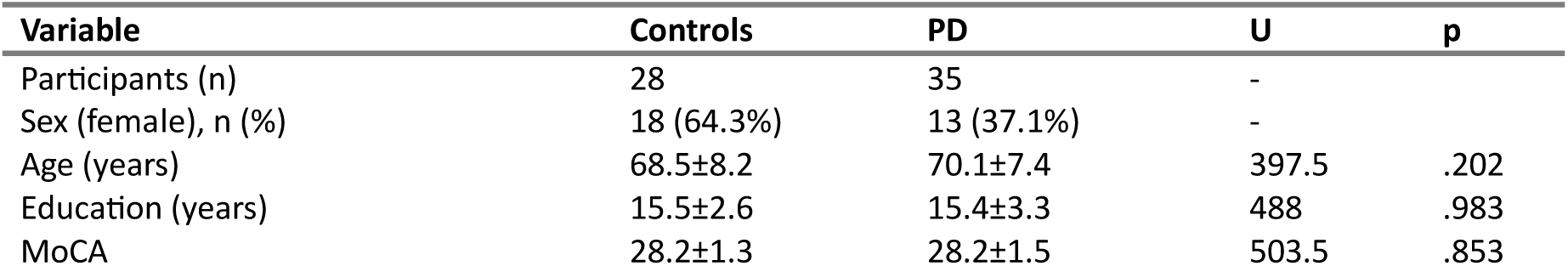
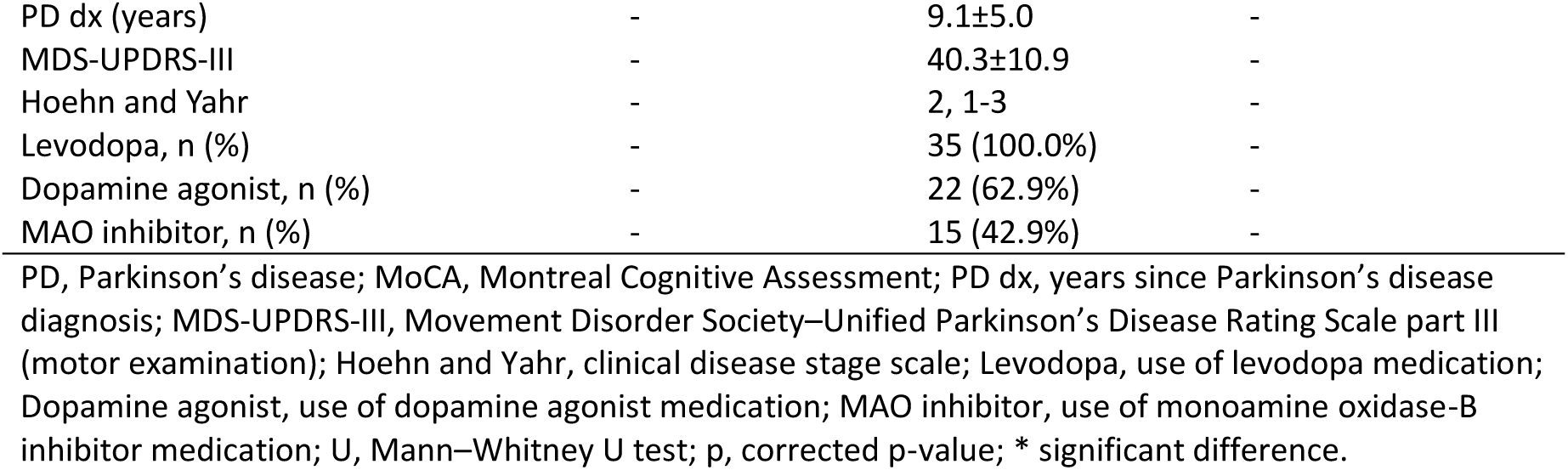
Demographic and behavioural questionnaires.

| Variable | Controls | PD | U | p |
| --- | --- | --- | --- | --- |
| Participants (n) | 28 | 35 | - |  |
| Sex (female), n (%) | 18 (64.3%) | 13 (37.1%) | - |  |
| Age (years) | 68.5±8.2 | 70.1±7.4 | 397.5 | .202 |
| Education (years) | 15.5±2.6 | 15.4±3.3 | 488 | .983 |
| MoCA | 28.2±1.3 | 28.2±1.5 | 503.5 | .853 |

|  |  |  |  |
| --- | --- | --- | --- |
| PD dx (years) | - | 9.1±5.0 | - |
| MDS-UPDRS-III | - | 40.3±10.9 | - |
| Hoehn and Yahr | - | 2, 1-3 | - |
| Levodopa, n (%) | - | 35 (100.0%) | - |
| Dopamine agonist, n (%) | - | 22 (62.9%) | - |
| MAO inhibitor, n (%) | - | 15 (42.9%) | - |

PD, Parkinson's disease; MoCA, Montreal Cognitive Assessment; PD dx, years since Parkinson's disease diagnosis; MDS-UPDRS-III, Movement Disorder Society–Unified Parkinson's Disease Rating Scale part III (motor examination); Hoehn and Yahr, clinical disease stage scale; Levodopa, use of levodopa medication; Dopamine agonist, use of dopamine agonist medication; MAO inhibitor, use of monoamine oxidase-B inhibitor medication; U, Mann–Whitney U test; p, corrected p-value; \* significant difference.

Participants with PD were adults with clinically probable PD according to Movement Disorder Society (MDS) criteria [28] diagnosed in outpatient neurology clinics around the country by certified neurologists. PD participants were excluded if their symptoms exceeded stage III on the Hoehn & Yahr scale [29], were in treatment with deep brain stimulation, and/or suffered from severe walking and balance problems. Participants were excluded if they had uncontrolled metabolic problems, hearing loss, or any other neurological or psychiatric disorders. Participants with PD were instructed to maintain their regular medication regimen which included levodopa for all participants.

Participants were recruited via posters, social media, and through the Danish Parkinson’s Association (Parkinsonsforeningen).

All participants provided written and verbal consent before the two-hour experiment. The study was approved by the Institutional Review Board at the DNC (Dansk Neuroforskningscenter), Aarhus University Hospital, Aarhus, Denmark (DNC-IRB-2021-010), and conducted in accordance with the ethical standards laid down in the 1964 Declaration of Helsinki and its later amendments. The study was not prospectively registered. Controls received 200 DKK in compensation for their participation. Participants with PD did not receive monetary compensation for their participation, instead, the costs of transportation were reimbursed.

### 2.2. Demographic and behavioural questionnaires

Participants answered a background questionnaire that included demographic data such as age, sex, education level, family medical history, and current PD medication. Then, the Montreal Cognitive Assessment (MoCA) was conducted, to exclude mild cognitive impairment (MCI) (cut-off=26). Finally, PD symptomatology was examined via the motor examination of the Movement Disorder Society - Unified Parkinson’s disease rating scale (MDS-UPDRS-III) to ensure clinically probable PD and assess the severity of the disease. These assessments were conducted by VPN.

### 2.3. Task and Stimuli

The synchronisation task was conducted using Max MSP (version 8.1.2). Tapping responses were recorded using an Apple Keyboard (Apple Inc., USA). To correct for input latency, all taps had 96 ms subtracted, which corresponded to combined hard- and software delay, using Audacity (version 3.4.2) and MATLAB (version R2023b). All participants completed a practice trial before the experimental task. During each trial, participants listened to a musical sequence and were instructed to tap along with the beat as accurately and consistently as possible for the full duration of the sequence. The stimuli comprised four 40-s music clips presented at different tempi: 100, 110, 120, and 130 beats per minute. The clips consisted of bass drum, snare drum, hi-hat, bass, and organ, with accents at the beat level and no syncopation. They were the low-complexity versions of the stimuli used in [30].

### 2.4. Data pre-processing

Data consisted of finger-tap times and pacing beat onset times estimated from each of the four stimulus audio files, which differed in tempo (100, 110, 120, and 130 BPM). Beat intervals in these stimulus files were 600 ms, ∼545 ms, 500 ms, or ∼462 ms.

For ADAM parameter estimation, matched pairs of finger-tap and beat-onset times are required for each experimental trial. In cases where participants produce one tap per beat (as instructed), these matched pairs are already in the raw data. However, in cases where participants miss taps or produce extra taps, data pre-processing is required to extract matched pairs. Before removal of outlier trials for each dependent measure (see ‘2.6. Data analysis’), the number of taps produced per trial ranged from 48 to 103 (median = 60) for PD participants and 49 to 108 (median = 79) for controls. After outlier removal, the number of taps produced per trial ranged from 48 to 92 (median = 59) for PD participants and 49 to 108 (median = 79) for controls.

Matched tap-beat pairs were extracted by stepping through each trial beat by beat (starting with the 9^th^ beat) and finding the nearest tap. Specifically, onset time differences between the stimulus beat onset and all tap times were computed at each step, and the beat-tap pair with smallest absolute onset time difference was selected. This resulted in 58 matched pairs per trial.

Missing taps and outlier taps were identified based on a criterion of 0.5 of the pacing sequence inter-onset interval. Onset time differences larger than this value were coded as ‘NaN’. Trials were then screened for missing data based on a criterion of five missing values (NaNs), of which there were no consecutive cases. Only trials that met these criteria were deemed complete and included in the analyses. Other trials were deemed incomplete and were not analysed. The number of trials before and after removing incomplete trials is shown in **Table 2**.

**Table 2:**
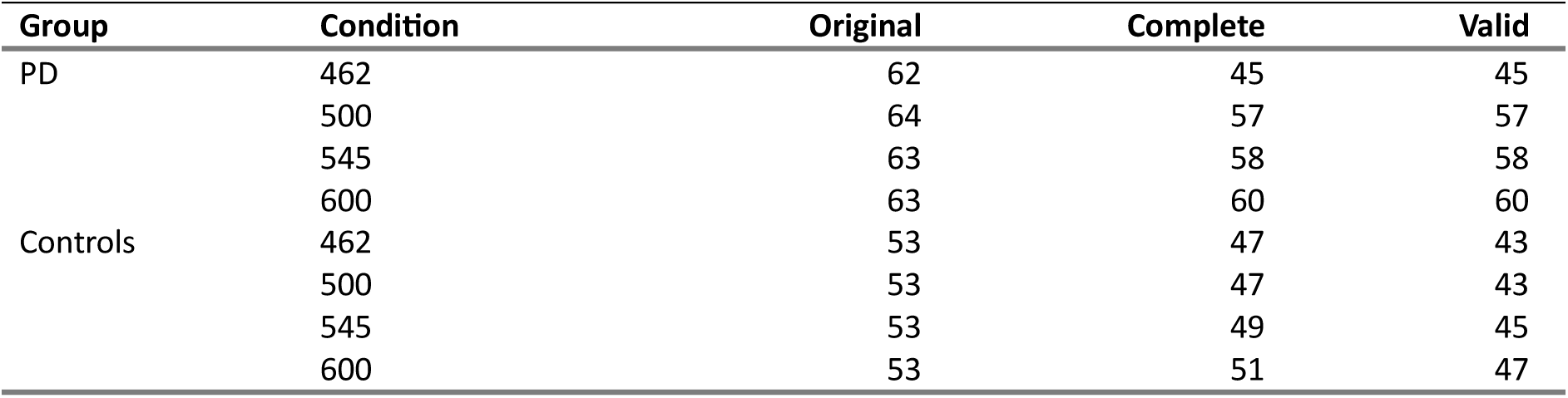
Number of trials produced in each participant Group and Tempo condition for the original data set (original), after screening for missing and outlier taps (complete), and after removing outlier trials (valid). PD = Parkinson’s Disease.

The gaps in valid trials were repaired via linear interpolation, as in previous studies [21, 27, 31]. In the final sample, interpolation was required in 27% of the total number of trials in the PD group (filling a total of 140 gaps, amounting to 0.96% of all target tap-beat pairs) and 4% of trials in the control group (filling a total of 11 gaps, amounting to 0.10% of all target tap-beat pairs). Note that the actual number of interpolated events was negligible (around 1%) in both groups.

### 2.5. Measures of synchronization performance and ADAM parameters

#### 2.5.1. Synchronization performance

The quality of sensorimotor synchronization performance was assessed by computing the temporal asynchrony between stimulus events and corresponding tap responses in each valid trial. Pacing event onset times were subtracted from tap times, resulting in negative asynchronies if taps preceded pacing sounds. Inter-stimulus (beat onset) intervals and inter-response (tap) intervals were also computed for use as input in the ADAM parameter estimation procedure.

Three main measures of synchronization were extracted from each trial: (1) Mean signed asynchrony, which serves as a measure of how early or late participants tap, on average, relative to beat onsets, (2) mean absolute asynchrony, which provides a global measure of synchronization accuracy, and (3) the variability (coefficient of variation) of signed asynchronies, which is an inverse index of synchronization stability. The coefficient of variation (CV) was calculated by dividing the standard deviation of the asynchronies by the mean inter-tap interval in each valid trial.

#### 2.5.2. ADAM Parameter estimation

ADAM parameter estimates were obtained for each valid trial in each Tempo condition (600, 545, 500, and 462 ms) for each participant in each Group (PD and controls).

ADAM was designed as a tool for exploring interactions between sensory-motor and cognitive processes for temporal adaptation, anticipation, and attention allocation, as postulated in the ‘ensemble skills’ theoretical framework . ADAM consists of three computational modules that are assumed to interact via internal models that represent sensory-motor transformations supporting action planning and prediction related to ‘self’ and ‘other’ [14–17, 20, 21] (**Figure 1**). Each process instantiated in ADAM is controlled by a separate parameter, and the value of these parameters can be estimated for a particular individual by fitting the model to time series data from sensorimotor synchronization tasks [21, 24, 27, 31].

ADAM’s *adaptation module* compensates for synchronization errors by implementing phase correction and period correction and determines the provisional timing of the next planned movement by providing input to an internal model of the ‘self’ [32]. In line with the two-level model of variability in movement timing [33, 34] and its extension to sensorimotor synchronization [3, 4, 35–37], ADAM includes two sources of noise, one that represents variability in a ‘timekeeper’ in the central nervous system and the other representing ‘motor’ noise in the peripheral nervous system.

Components of ADAM and the underlying equations are shown in **Figure 2**. The main model parameters are phase correction (α) and period correction (β) from ADAM’s adaptation module, temporal prediction/tracking (δ) from the anticipation module, and anticipatory error correction (γ) from the joint module. Parameters are also estimated for central ‘timekeeper’ noise (Tn) and peripheral ‘motor’ noise (Mn). The estimation procedure used the bounded Generalised Least Squares method [2, 23], which harnesses the analytic efficiency of matrix algebra applied to behavioural time series data [22].

ADAM parameters were estimated for each participant using MATLAB scripts adapted from [21]. Data from each valid trial were transformed into the appropriate structure, which consists of three time series of equal length. The modelling procedure treats one series as reference time intervals associated with a pacing sequence (‘inter-stimulus intervals’), another series as time intervals associated with movements produced by an individual synchronizing with the pacing sequence (‘inter-response intervals’), and a final series consisting of the ‘asynchronies’ between the onsets of corresponding events in the pacing sequence and the response sequence (again, stimulus onset times are subtracted from response onset times, yielding positive asynchronies if responses occur later than stimuli).

Parameter estimates can be obtained, in principle, using adaptation-only and full ‘joint’ versions of ADAM [27]. The adaptation-only version includes phase correction (α), period correction (β), timekeeper noise (*Tn*), and motor noise (*Mn*). In the current study, which required sensorimotor synchronization with pacing sequences at a steady tempo, parameters estimates were only obtained from the adaptation-only version of ADAM. Previous studies with computer-controlled pacing sequences and computer simulations [20, 21, 27] suggest that temporal adaptation is sufficient to account for sensorimotor synchronization at steady tempi.

### 2.6. Data analysis

Statistical analyses were performed using R (version 4.3.1) running in RStudio (version 2023.09.1+494). Outlier trials were identified for the main behavioural outcome dependent variables (mean absolute asynchrony and CV of asynchronies) within each Group × Tempo cell using the interquartile range (IQR) method as implemented in rstatix (version 0.7.2). Observations exceeding 1.5 × IQR were flagged as mild outliers, and those exceeding 3 × IQR were classified as extreme outliers. Data from participants with extreme outliers were removed from subsequent analyses. Outlier removal resulted in valid datasets for n = 35 PD participants (0 were removed) and n = 28 controls (2 were removed). The number of valid trials following outlier removal are shown in **Table 2**.

Shapiro-Wilk tests were used to check whether data for each dependent variable (asynchrony measures and ADAM parameter estimates) met the assumption of normal distribution using the ‘shapiro_test’ function from the ‘rstatix’ R package (version 0.7.2). The ‘bestNormalize’ package (version 1.9.1) was used to select the most appropriate normalization method for each variable, when required. Where substantial deviations from normality were observed (operationalized as a majority of Group × Tempo cells yielding significant Shapiro–Wilk tests), data transformations were applied. Log-transformation was used for mean absolute asynchrony and the CV of asynchronies. Ordered-quantile (ORQ) rank-based inverse normal transformation (‘orderNorm’) was used for other variables (mean signed asynchrony and ADAM parameters), which could have zero or negative values. Transformed variables were used as dependent variables in subsequent modelling.

Given that linear mixed-effects models (LMMs) assume normally distributed residuals rather than raw scores, additional diagnostic checks were conducted at the model level. For each dependent variable, a preliminary LMM was fit with fixed effects of Group, Tempo, and their interaction, and a random intercept for Participant. Residuals from these models were extracted and subjected to Shapiro–Wilk tests to evaluate adherence to normality assumptions. In addition, visual inspection of data distributions was conducted for each dependent variable. Histograms, quantile–quantile (Q–Q) plots, and boxplots were generated as a function of participant Group and Tempo condition. These visualizations facilitated assessment of distributional shape, variance homogeneity, and the presence of outliers, and were used to corroborate the results of formal statistical tests.

Three main sets of analyses were conducted. The first set tested effects of participant group (‘Group’) and tempo condition (‘Tempo’) on sensorimotor synchronization performance by examining each behavioural outcome variable separately: mean signed asynchrony, mean absolute asynchrony, and the CV of asynchronies. The second set tested effects of Group and Tempo on estimates of each ADAM parameter: Phase correction, period correction, timekeeper noise, and motor noise. The third set of analysis aimed to identify the set of ADAM parameter estimates that best accounted for synchronization performance across the different participant Groups and Tempo conditions. These analyses thus included ADAM parameter estimates, along with Group and Tempo, as predictors of each behavioural outcome dependent variable. The third set included interactions between each ADAM parameter and Group, but did not include interactions with Tempo, which was included as a separate fixed factor. A fourth set of analyses included interactions between each ADAM parameter, Group, and Tempo, and it is only reported in the **Supplementary Information**. All dependent variables for the four sets of analyses were screened for outliers and tested for normality.

In the first two sets of analyses, separate linear mixed-effects analyses were conducted for each dependent measure. The main mixed-effects models, which included participant Group and Tempo condition as fixed factors and participant as a random effect variable, were compared with reduced models, containing only the random effects, using the ‘anova’ R function. The log-likelihood, the Akaike Information Criterion (AIC), and the Bayesian Information Criterion (BIC) were used to assess model fit. For each mixed model, main effects and interactions of Tempo condition and participant Group were tested using the ‘joint_tests’ function from the ‘emmeans’ R package, and planned orthogonal contrasts tested linear, quadratic, and cubic polynomial trends across the four Tempo conditions using the ‘contrast’ function from the ‘lsmeans’ R package. The full set of contrasts is shown in **Table S4**. The linear contrast tests hypotheses about changes in dependent measures with decreasing Tempo, while the quadratic contrast tests hypotheses about differences for intermediate Tempo (i.e., 500 ms and 545 ms inter-beat intervals), which are close to typical preferred Tempo, versus faster and slower Tempo. The cubic trend was included to test whether these linear and quadratic components fully capture the effects-of-interest or whether greater complexity is present in the data patterns.

In the third set of analyses, separate linear mixed effects models for each asynchrony dependent measure included fixed factors of phase correction, period correction, timekeeper noise, and motor noise, their two-way interactions with Group, Tempo as a covariate (which was not included in interactions), as well as participant as a random effects variable. The formula for the linear mixed effect model in this set is as follows:

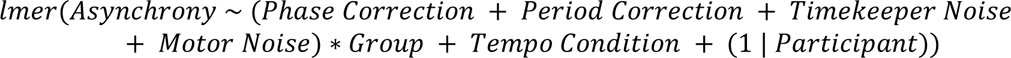

To check for multicollinearity between ADAM parameter estimates, we calculated variance inflation factors (VIFs) using the ‘vif.mer’ function (adapted from https://github.com/aufrank/R-hacks/blob/master/mer-utils.R). This check was done in versions of the linear mixed-effects models that omitted interactions with “Group” and “Condition”. VIFs greater than 5 were considered potentially problematic [38] but all observed values were below this criterion. The main mixed-effects models were compared with a reduced model containing only random effects. Statistical interactions in the main mixed model were unpacked with simple slopes analyses using the ‘probe_interaction’ function from the ‘interactions’ R package (version 1.1.5).

## 3. Results

### 3.1. Synchronization performance

Data for mean signed asynchrony, mean absolute asynchrony, and the variability (CV) of asynchronies for the PD and control group across the four Tempo conditions are plotted in **Figure 3**.

**Figure 3:**
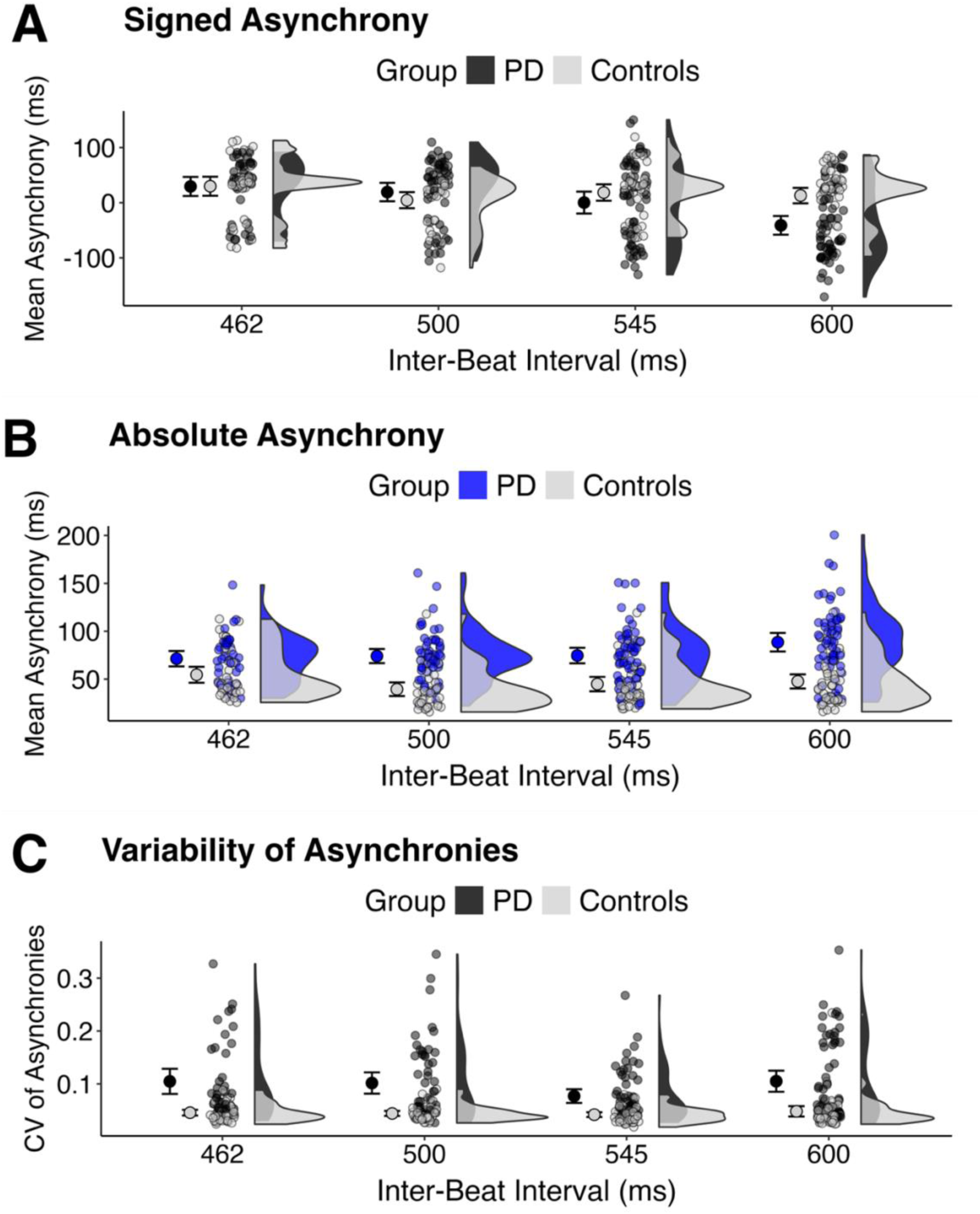
Measures of synchronization performance for participants in the PD and control groups in the four Tempo conditions (462, 500, 545, and 600 ms inter-beat intervals): (A) mean signed asynchrony (ms); (B) mean absolute asynchrony (ms); (C) variability of asynchronies (CV). Note that all asynchrony data were transformed for analysis, but untransformed values are plotted here. PD = Parkinson’s Disease.

For mean signed asynchrony (which were ORQ transformed in the analysis), model comparison revealed that the main mixed-effects model fit the data better than the reduced linear model that included only random effects (χ2 (7) = 164.75, *p* < 0.001; Log Likelihood = -305.74 (main) vs - 388.12 (reduced)). The main effect of Tempo condition was statistically significant, *F*(3, 317.08) = 49.03, *p* < .001, as was the interaction between Group and Tempo, *F*(3, 317.08) = 15.53 *p* < .001. The main effect of Group was not significant (*p* = .313). The polynomial contrast analysis revealed statistically significant linear and cubic trends across Tempo, and significant interactions of Group with the linear and quadratic trends across Tempo (**Table 3**). No other effects were statistically significant. The linear trend indicates that mean signed asynchrony went from being positive (i.e., taps occurred after beats) at the fastest Tempo (462 ms inter-beat intervals) to being negative (taps preceded beats) at the slowest Tempo (600 ms) (Figure 3A). The interaction with Group indicates that this effect was more pronounced for PD than controls. The interaction of Group with the quadratic trend likely reflects a U-shaped function only in controls, where asynchronies were closest to zero at intermediate Tempo (500 ms and 545 ms) but otherwise were more positive. The interpretation of the cubic trend is not straightforward.

**Table 3:** Outcomes of the contrast analysis for mean signed asynchrony. [Confidence intervals were also computed but are not displayed.] *** p < 0.001; ** p < 0.01; * p < 0.05.

| Effect | Contrast | Estimate | SE | Df | t value | p value |
| --- | --- | --- | --- | --- | --- | --- |
| Group | PD vs Controls | 0.988 | 0.971 | 61.354 | 1.017 | 0.313 |
| Tempo | Linear | -4.611 | 0.418 | 318.390 | -11.036 | <0.001*** |
|  | Quadratic | 0.243 | 0.186 | 317.804 | 1.309 | 0.191 |
|  | Cubic | -1.825 | 0.412 | 317.089 | -4.434 | <0.001*** |
| Group x Tempo | Group x Linear | 2.223 | 0.418 | 318.390 | 5.320 | <0.001*** |
|  | Group x Quadratic | -0.660 | 0.186 | 317.804 | -3.555 | <0.001*** |
|  | Group x Cubic | -0.691 | 0.412 | 317.089 | -1.680 | 0.094 |

For mean absolute asynchrony data, which were log-transformed, the main mixed-effects model fit the data better than the reduced linear model that included only random effects (χ2 (7) = 56.379, *p* < .001; Log Likelihood = -119.52 (main) vs -147.71 (reduced)). There were statistically significant main effects of Group, *F*(1, 61.30) = 20.57, *p* < .0001, and Tempo, *F*(3, 318.46) = 5.52, *p* = .001, and a significant Group x Tempo interaction, *F*(3, 318.46) = 7.92, *p* < .001. The main effect of Group reflected higher absolute asynchrony in PD than controls. The polynomial contrast analysis (**Table 4**) revealed the effect of Tempo was driven by a quadratic trend, where absolute asynchrony was, on average, lower at intermediate Tempo than at fast or slow Tempo. The contrasts also indicated that the Group x Tempo interaction had two components. First, a linear trend, whereby asynchrony increased as Tempo got slower, was evident in PD participants but not controls. By contrast, visual inspection of Figure 3B reveals that the pattern displayed by controls appears like a mix of quadratic and cubic function, perhaps driving the interaction between Group and the cubic trend for Tempo. No other effects were statistically significant.

**Table 4:** Outcomes of the contrast analysis for mean absolute asynchrony. *** p < 0.001; ** p < 0.01; * p < 0.05.

| Effect | Contrast | Estimate | SE | Df | t value | p value |
| --- | --- | --- | --- | --- | --- | --- |
| Group | PD vs Controls | -1.855 | 0.409 | 61.302 | -4.535 | <0.001*** |
| Tempo | Linear | -0.185 | 0.270 | 320.554 | -0.686 | 0.493 |
|  | Quadratic | -0.464 | 0.120 | 319.593 | -3.866 | <0.001*** |
|  | Cubic | -0.382 | 0.266 | 318.278 | -1.437 | 0.152 |
| Group x Tempo | Group x Linear | -0.966 | 0.270 | 320.554 | -3.584 | <0.001*** |
|  | Group x Quadratic | -0.196 | 0.120 | 319.593 | -1.631 | 0.104 |
|  | Group x Cubic | -0.828 | 0.266 | 318.278 | -3.114 | 0.002** |

For CV of asynchronies, which were log-transformed, the main mixed-effects model fit the data better than the reduced linear model (χ2 (7) = 31.09, *p* < .001; Log Likelihood = -108.15 (main) vs - 123.69 (reduced)). Statistically significant main effects were observed for Group, *F*(1, 61.4) = 14.94, *p* < .001, and Tempo, *F*(3, 317.1) = 3.635, *p* = .013, but Group x Tempo interaction was not significant (*p* > .05). The Group main effect reflected higher CV of asynchronies in PD than controls (**Figure 3c**). The polynomial contrast analysis revealed statistically significant quadratic and cubic trends across Tempo (**Table 5**). Variability was lower for intermediate Tempo than fast or slow Tempo but was also slightly higher for intermediate 500 ms than 545 ms inter-beat intervals. No other effects were statistically significant.

**Table 5:** Outcomes of the contrast analysis for the variability of asynchronies. *** p < 0.001; ** p < 0.01; * p < 0.05.

| Effect | Contrast | Estimate | SE | Df | t value | p value |
| --- | --- | --- | --- | --- | --- | --- |
| Group | PD vs Controls | -2.143 | 0.554 | 61.400 | -3.865 | <0.001*** |
| Tempo | Linear | -0.245 | 0.246 | 318.469 | -0.997 | 0.319 |
|  | Quadratic | -0.251 | 0.109 | 317.857 | -2.297 | 0.022* |
|  | Cubic | 0.507 | 0.242 | 317.108 | 2.093 | 0.037* |
| Group x Tempo | Group x Linear | -0.172 | 0.246 | 318.469 | -0.700 | 0.485 |
|  | Group x Quadratic | 0.162 | 0.109 | 317.857 | 1.480 | 0.140 |
|  | Group x Cubic | -0.288 | 0.242 | 317.108 | -1.191 | 0.235 |

### 3.2. ADAM parameter estimates

Data for ADAM parameter estimates of phase correction, period correction, timekeeper noise, and motor noise for the PD and control group across the four Tempo conditions are plotted in **Figure 4**. Results of analyses for each parameter estimate are reported in separate sections below.

**Figure 4:**
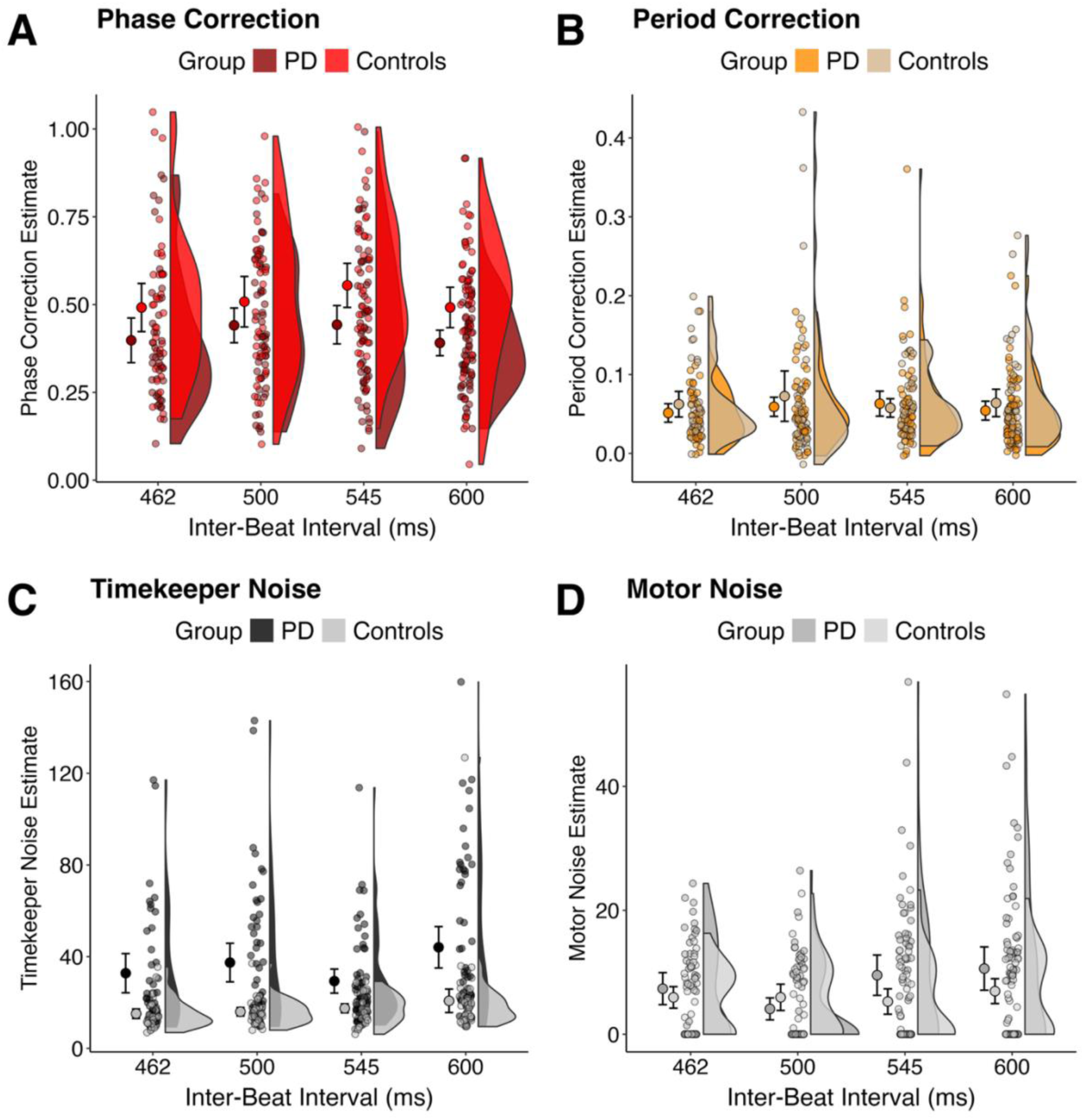
ADAM parameter estimates for participants in the PD and control groups in the four Tempo conditions (462, 500, 545, and 600 ms inter-beat intervals): (A) phase correction; (B) period correction; (C) timekeeper noise; (D) motor noise. Raw data are plotted, but data were ordered quantile (ORQ) transformed prior to analysis. PD = Parkinson’s Disease.

#### 3.2.1. Phase correction

The main linear mixed-effects model for phase correction estimates (ORQ-transformed) fit the data better than the reduced linear model that included only random effects (χ2 (7) = 16.857, *p = .*018; Log Likelihood = -504.67 (main) vs -513.10 (reduced)). A statistically significant main effect was observed for Group, *F*(1, 57.76) = 11.41, *p* = .001, but the effect of Tempo and the Group x Tempo interaction were not statistically significant, *p*s > .05. The main effect of Group reflects lower phase correction estimates for PD than controls (**Figure 4a**). The polynomial contrast analysis revealed a significant quadratic across Tempo (**Table 6**). Phase correction estimates were higher at intermediate Tempo (500 ms and 545 ms) than at fast (462 ms) and slow (600 ms) Tempo for PD and controls.

**Table 6:** Outcomes of the contrast analysis for ADAM phase correction estimates. *** p < 0.001; ** p < 0.01; * p < 0.05.

| Effect | Contrast | Estimate | SE | Df | *t* value | *p* value |
| --- | --- | --- | --- | --- | --- | --- |
| Group | PD vs Controls | 1.839 | 0.544 | 57.764 | 3.378 | 0.001** |
| Tempo | Linear | 0.013 | 0.873 | 330.199 | 0.015 | 0.988 |
|  | Quadratic | 0.839 | 0.389 | 328.667 | 2.159 | 0.032* |
|  | Cubic | -0.676 | 0.865 | 325.361 | -0.782 | 0.435 |
| Group x Tempo | Group x Linear | 0.156 | 0.873 | 330.199 | 0.179 | 0.858 |
|  | Group x Quadratic | -0.054 | 0.389 | 328.667 | -0.140 | 0.889 |
|  | Group x Cubic | -0.824 | 0.865 | 325.361 | -0.953 | 0.341 |

#### 3.2.2. Period correction

The main mixed-effects model for period correction (ORQ-transformed) did not fit the data better than the reduced linear model (χ2 (7) = 2.511, *p = .*926; Log Likelihood = -520.66 (main) vs -521.91 (reduced). Indeed, the main effects of Group and Tempo, and the Group x Tempo interaction, were all not statistically significant (*p*s > .05) (**Figure 4b**). The polynomial contrast analysis likewise did not reveal any significant effects (**Table 7**). Note that this lack of reliable effects of Group and Tempo on period correction could be expected because the pacing sequences did not contain tempo changes.

**Table 7:** Outcomes of the contrast analysis for ADAM period correction estimates. *** p < 0.001; ** p < 0.01; * p < 0.05.

| Effect | Contrast | Estimate | SE | Df | *t* value | *p* value |
| --- | --- | --- | --- | --- | --- | --- |
| Group | PD vs Controls | 0.378 | 0.491 | 56.533 | 0.770 | 0.445 |
| Tempo | Linear | 0.116 | 0.932 | 333.165 | 0.124 | 0.901 |
|  | Quadratic | 0.163 | 0.415 | 331.186 | 0.393 | 0.694 |
|  | Cubic | -0.610 | 0.925 | 327.396 | -0.660 | 0.510 |
| Group x Tempo | Group x Linear | -0.095 | 0.932 | 333.165 | -0.102 | 0.919 |
|  | Group x Quadratic | -0.438 | 0.415 | 331.186 | -1.055 | 0.292 |
|  | Group x Cubic | -0.296 | 0.925 | 327.396 | -0.320 | 0.749 |

#### 3.2.3. Timekeeper noise

The main linear mixed-effects model for timekeeper noise estimates (ORQ-transformed) fit the data better than the reduced linear model (χ2 (7) = 47.48, *p* < .001; Log Likelihood = -382.65 (main) vs - 406.39 (reduced). There were significant main effects of Group, *F*(1, 61.29) = 12.32, *p* < .001, and Tempo, *F*(3, 318.54) = 11.80, *p* < .001, but the Group x Tempo interaction was not statistically significant (*p* > .05). The effect of Group indicated that timekeeper noise estimates were higher in PD than controls (**Figure 4c**), while the Tempo effect reflects increasing timekeeper noise estimates with decreasing Tempo (**Table 8**).

**Table 8:** Outcomes of the contrast analysis for ADAM timekeeper noise estimates. *** p < 0.001; ** p < 0.01; * p < 0.05.

| Effect | Contrast | Estimate | SE | Df | *t* value | *p* value |
| --- | --- | --- | --- | --- | --- | --- |
| Group | PD vs Controls | -2.889 | 0.823 | 61.287 | -3.510 | 0.001** |
| Tempo | Linear | 3.236 | 0.551 | 320.660 | 5.867 | <0.001*** |
|  | Quadratic | -0.030 | 0.245 | 319.687 | -0.124 | 0.901 |
|  | Cubic | 0.399 | 0.544 | 318.345 | 0.734 | 0.464 |
| Group x Tempo | Group x Linear | 0.042 | 0.551 | 320.660 | 0.076 | 0.940 |
|  | Group x Quadratic | 0.207 | 0.245 | 319.687 | 0.845 | 0.399 |
|  | Group x Cubic | -0.359 | 0.544 | 318.345 | -0.660 | 0.510 |

#### 3.2.4. Motor noise

For motor noise estimates (ORQ-transformed), model comparison revealed that the main mixed-effects model fit the data better than the reduced model (χ2 (7) = 15.998, *p* < 0.05; Log Likelihood = - 456.6 (main) vs -464.6 (reduced)). However, the main effects and interaction of Group and Tempo were all not statistically significant (ps > .05) (**Figure 4d**). The polynomial contrast analysis revealed a statistically significant interaction between Group and the cubic trend across Tempo (**Table 9**).

**Table 9:** Outcomes of the contrast analysis for ADAM motor noise estimates. *** p < 0.001; ** p < 0.01; * p < 0.05.

| Effect | Contrast | Estimate | SE | Df | t value | p value |
| --- | --- | --- | --- | --- | --- | --- |
| Group | PD vs Controls | -0.414 | 0.456 | 57.188 | -0.908 | 0.368 |
| Tempo | Linear | 1.200 | 0.772 | 331.285 | 1.555 | 0.121 |
|  | Quadratic | -0.641 | 0.344 | 329.640 | -1.864 | 0.063 |
|  | Cubic | -0.942 | 0.765 | 326.178 | -1.231 | 0.219 |
| Group x Tempo | Group x Linear | -0.645 | 0.772 | 331.285 | -0.836 | 0.404 |
|  | Group x Quadratic | 0.234 | 0.344 | 329.640 | 0.680 | 0.497 |
|  | Group x Cubic | 1.577 | 0.765 | 326.178 | 2.061 | 0.040* |

### 3.3. Relations between synchronization performance and ADAM parameter estimates

This analysis aimed to identify the set of ADAM parameter estimates that best accounted for synchronization performance in the PD and control groups. Here, we included interactions between each ADAM parameter and Group, but did not include interactions with Tempo, which was included as a separate fixed factor and reported only in the **Supplementary Information**.

Relations between mean signed asynchrony and each ADAM parameter are shown for each participant group in **Figure 5**. The linear mixed effects model on these data included fixed factors of phase correction, period correction, timekeeper noise, and motor noise, their interactions with Group, and Tempo as a separate covariate, as well as participant as a random effects variable. Model comparison revealed that this main mixed-effects model fit the data significantly better than a reduced model with only random effects (χ2 (12) = 139.17, *p < .*001; Log Likelihood = -318.53 (main) vs -388.12 (reduced). In the main model (**Table 10**), period correction predicted signed asynchrony, with higher period correction associated with more positive asynchrony. There was also an effect of Tempo, corresponding to the effect described above in section 3.1, where it was seen that asynchrony becomes less positive/more negative as Tempo decreases. All other effects were non-significant.

**Figure 5:**
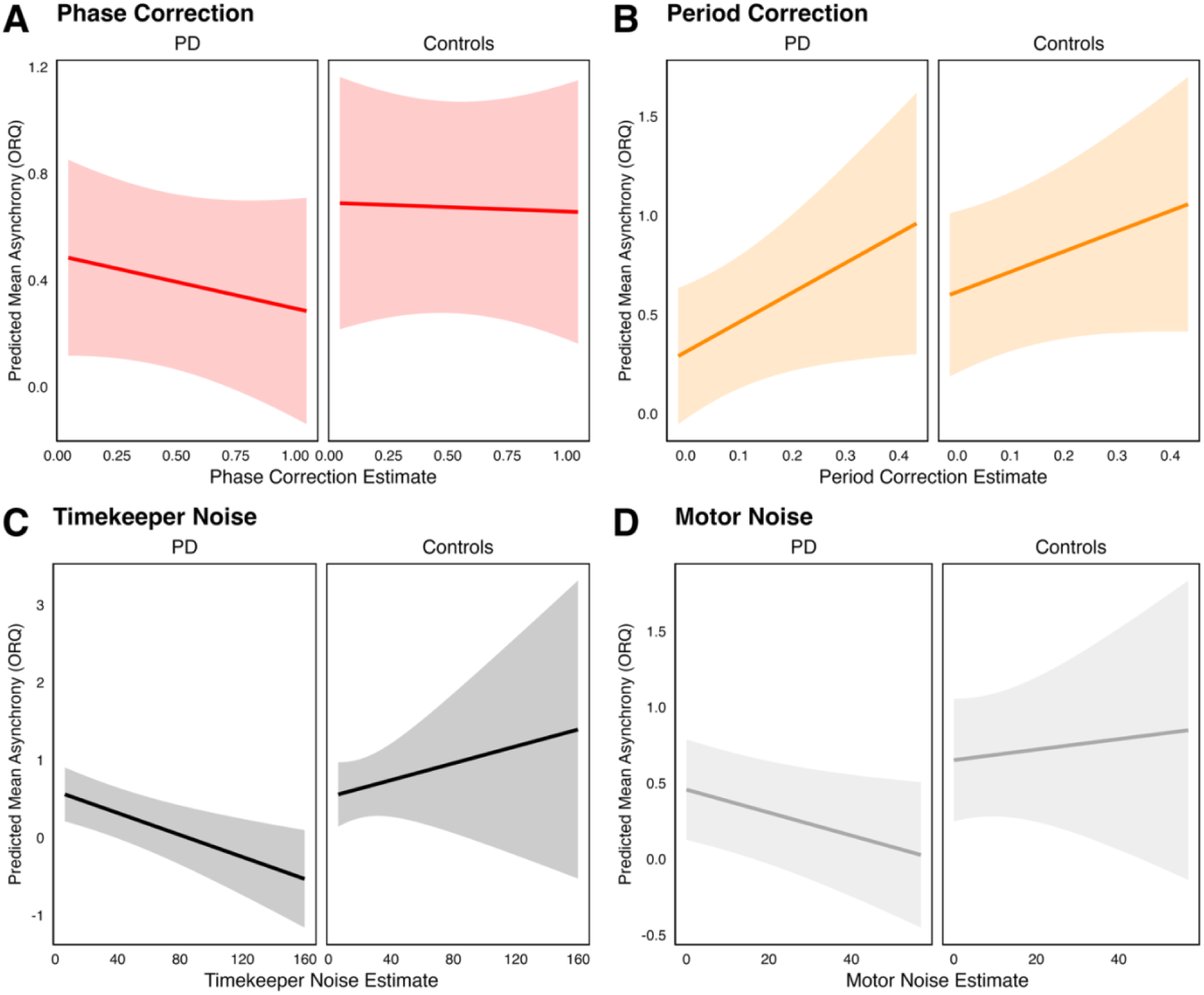
Simple slopes of relations between mean signed asynchrony (ORQ-transformed) and each ADAM parameter estimate in each participant group (collapsed across Tempo conditions). (A) Phase correction; (B) period correction; (C) timekeeper noise; (D) motor noise. PD = Parkinson’s Disease.

**Table 10:**
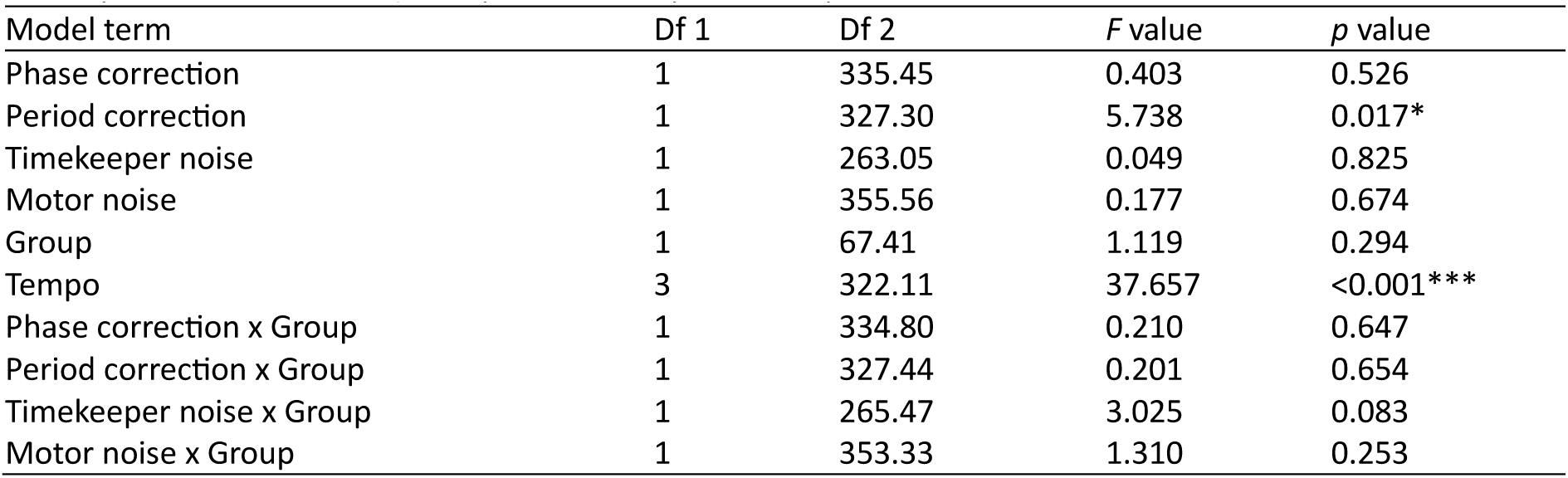
Effects of ADAM parameters, Group, and Tempo on mean signed asynchrony (excluding interactions of Tempo with other factors). *** p < 0.001; ** p < 0.01; * p < 0.05.

Relations between mean absolute asynchrony and each ADAM parameter are displayed in **Figure 6**. Model comparison revealed that the main mixed-effects model fit the data significantly better than a reduced random-effects-only model (χ2 (12) = 110.68, *p < .*001; Log Likelihood = -92.37 (main) vs - 147.71 (reduced). In the main model (**Table 11**), timekeeper noise estimates predicted absolute asynchrony, with higher timekeeper noise associated with larger mean asynchrony. Furthermore, there were main effects of Group and Tempo on asynchrony, corresponding to the effects described in section 3.1, where asynchrony was larger for PD than controls, and for fast and slow Tempo than intermediate Tempo (on average). All other effects were non-significant.

**Figure 6:**
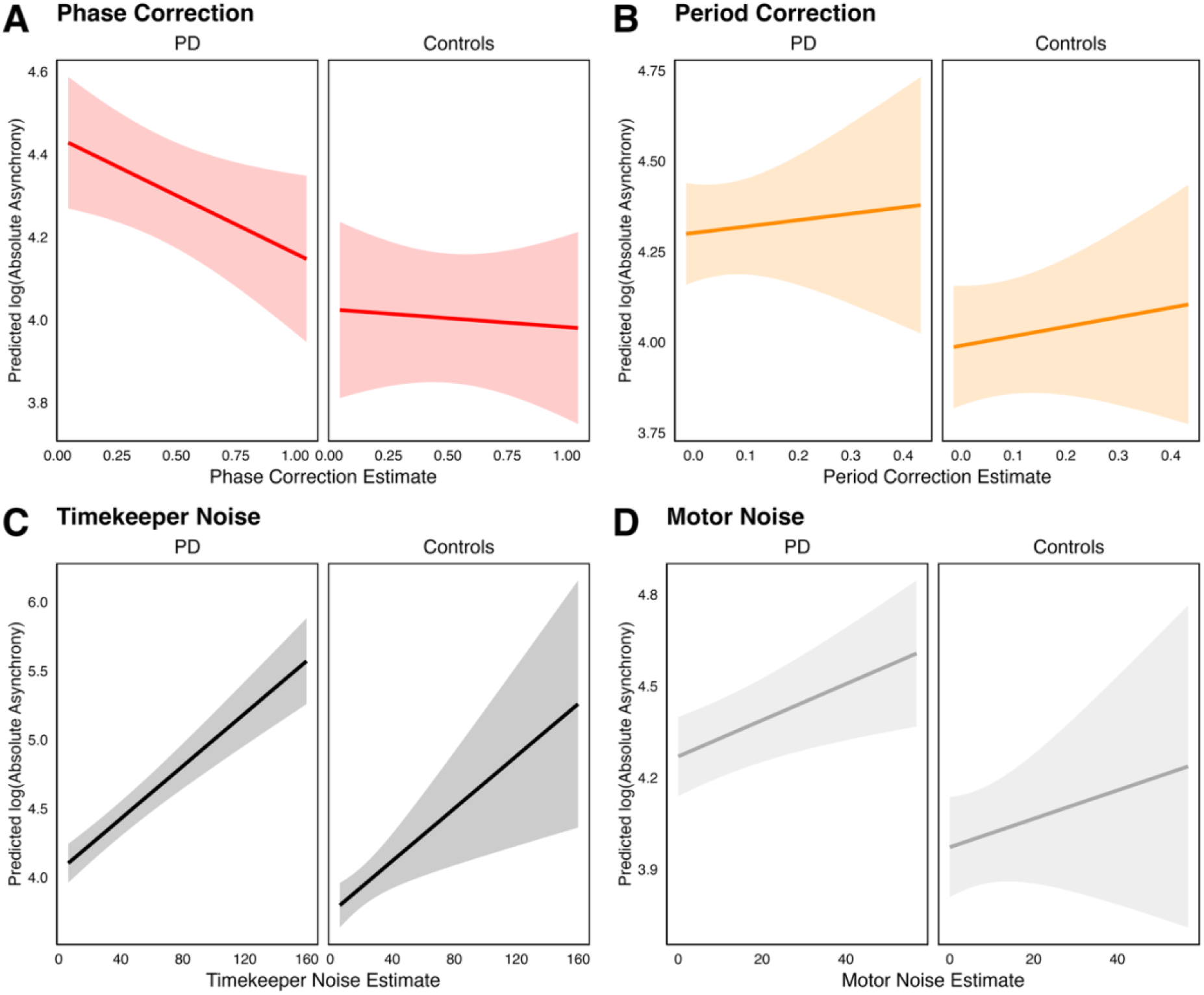
Simple slopes of relations between mean absolute asynchrony (log-transformed) and each ADAM parameter estimate in each participant group (collapsed across Tempo conditions). (A) Phase correction; (B) period correction; (C) timekeeper noise; (D) motor noise. PD = Parkinson’s Disease.

**Table 11:** Effects of ADAM parameters, Group, and Tempo on mean absolute asynchrony (excluding interactions of Tempo with other factors). *** p < 0.001; ** p < 0.01; * p < 0.05.

| Model term | Df 1 | Df 2 | F value | p value |
| --- | --- | --- | --- | --- |
| Phase correction | 1 | 351.12 | 2.367 | 0.125 |
| Period correction | 1 | 335.65 | 0.522 | 0.470 |
| Timekeeper noise | 1 | 230.11 | 29.365 | <0.001*** |
| Motor noise | 1 | 364.45 | 3.670 | 0.056 |
| Group | 1 | 73.21 | 9.970 | 0.002** |
| Tempo | 3 | 323.48 | 6.010 | 0.001** |
| Phase correction x Group | 1 | 350.56 | 1.287 | 0.257 |
| Period correction x Group | 1 | 335.76 | 0.019 | 0.889 |
| Timekeeper noise x Group | 1 | 235.83 | <0.001 | 0.991 |
| Motor noise x Group | 1 | 362.73 | 0.055 | 0.815 |

Relations between the CV of asynchronies and each ADAM parameter are shown in **Figure 7**. Model comparison revealed that the main mixed-effects model fit the data significantly better than a reduced random-effects-only model (χ2 (12) = 541.64, *p < .*001; Log Likelihood = 147.13 (main) vs - 123.69 (reduced). In the main model (**Table 12**), phase correction, timekeeper noise, and motor noise emerged as predictors of the CV of asynchronies, each individually and in interactions with Group. There were also main effects of Group and Tempo on the CV of asynchronies, as described in section 3.1, reflecting higher variability of asynchronies for PD than controls, and (non-linear) modulation of CV by Tempo. Regarding the effects for ADAM parameters, the CV of asynchronies decreased with increasing phase correction and decreasing timekeeper noise and motor noise. The pattern is consistent with the results of previous work with ADAM [24, 27]. The interactions with Group indicate that the dependency between phase correction and CV is stronger for PD than controls, whereas dependencies between timekeeper and motor noise parameters and CV are stronger for controls than PD participants. Note that interpreting these effects should consider that phase correction was generally lower, while timekeeper and motor noise were higher, for PD than controls.

**Figure 7:**
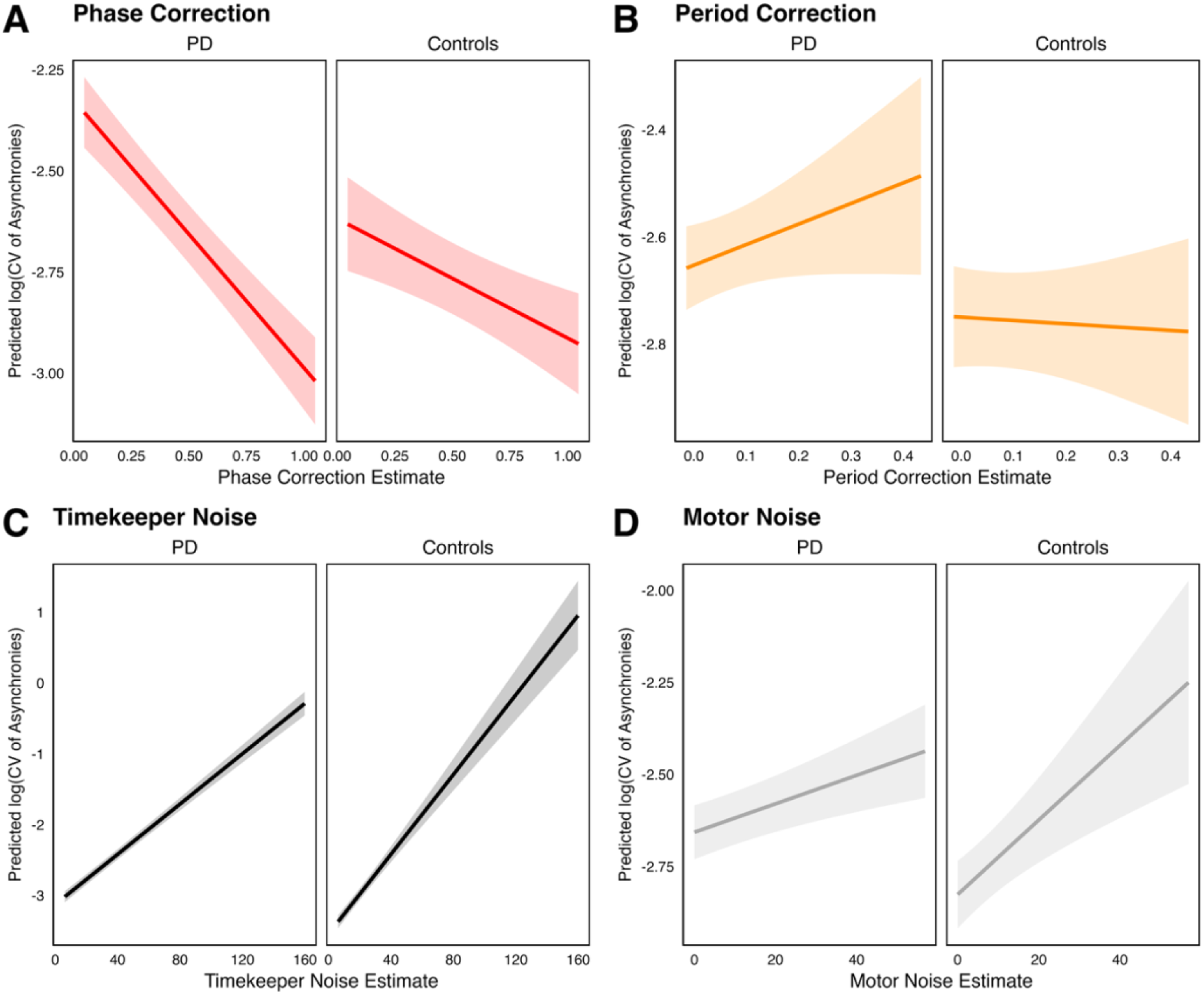
Simple slopes of relations between the CV of asynchronies (log-transformed) and each ADAM parameter estimate in each participant group (collapsed across Tempo conditions). (A) Phase correction; (B) period correction; (C) timekeeper noise; (D) motor noise. PD = Parkinson’s Disease.

**Table 12:** Effects of ADAM parameters, Group, and Tempo on the CV of asynchronies (excluding interactions of Tempo with other factors). *** p < 0.001; ** p < 0.01; * p < 0.05.

| Model term | Df 1 | Df 2 | *F* value | *p* value |
| --- | --- | --- | --- | --- |
| Phase correction | 1 | 344.90 | 78.175 | <0.001*** |
| Period correction | 1 | 331.65 | 1.065 | 0.303 |
| Timekeeper noise | 1 | 233.47 | 577.132 | <0.001*** |
| Motor noise | 1 | 360.45 | 24.071 | <0.001*** |
| Group | 1 | 72.55 | 4.934 | 0.029* |
| Tempo | 3 | 321.85 | 37.890 | <0.001*** |
| Phase correction x Group | 1 | 344.27 | 11.659 | 0.001** |
| Period correction x Group | 1 | 331.77 | 2.016 | 0.157 |
| Timekeeper noise x Group | 1 | 238.05 | 31.422 | <0.001*** |
| Motor noise x Group | 1 | 358.53 | 4.925 | 0.027* |

## 4. Discussion

The present study used the Adaptation and Anticipation Model (ADAM) to characterize the computational components of sensorimotor synchronization in Parkinson’s disease (PD). Three main findings emerged in the present study. First, participants with PD showed poorer synchronization performance than healthy controls, with larger absolute asynchronies and higher variability of asynchronies. Second, ADAM parameter estimates indicated lower phase correction and higher central timekeeper noise in PD, whereas period correction and motor noise did not show robust global group differences. Third, synchronization accuracy and stability were explained by partially distinct model parameters: timekeeper noise was the clearest predictor of absolute asynchrony, while phase correction, timekeeper noise, and motor noise jointly predicted synchronization variability, with group-dependent relations. Together, these results suggest that impaired rhythmic synchronization in PD reflects both reduced adaptive error correction and reduced precision of internal timing.

The main contribution of the study is therefore not simply to show that participants with PD synchronize tapping movements with rhythmic auditory cues less accurately. Rather, the results identify which timing components are most strongly associated with the impairment. Participants with PD were able to tap along with the auditory sequences, but their taps were, on average, farther from the beat and more variable from one beat to the next. In cognitive terms, the model suggests that this pattern was mainly related to two processes: participants with PD made weaker adjustments after tapping too early or too late, and their internal representation of the beat was less stable. Thus, the synchronization difficulties observed in PD appear to reflect altered error-based correction and internal timing precision, rather than a general inability to follow an external rhythm.

The absence of a robust global increase in motor noise suggests that the group difference was not merely an expression of variable finger movement execution. In other words, participants with PD did not simply tap more variably because each individual finger movement was noisier. Rather, the results point to difficulties in maintaining a stable internal beat and in using recent timing errors to adjust the next movement. This distinction is relevant for interpreting rhythmic auditory stimulation (RAS), because RAS can improve gait and other outcomes in PD, but its effects vary across individuals [6, 7, 39]. One possible reason for this variability is that benefit likely depends on the integrity of the timing mechanisms that allow an individual to trigger repetitive movements to couple these movements to the external cues via error correction [7, 12].

## Sensorimotor synchronization impairments in PD

At the behavioural level, the PD group showed larger absolute asynchronies and greater variability of asynchronies. These measures capture complementary aspects of synchronization. Absolute asynchrony indexes the magnitude of the temporal error regardless of whether taps occur early or late, whereas variability of asynchronies indexes how consistently the participant maintains alignment with the beat. The impairment in PD was therefore not best described as a simple directional bias in signed asynchrony (i.e., being consistently ahead of or lagging behind the pacing sequence). Instead, it reflected reduced precision and stability of auditory-motor alignment. This interpretation fits previous evidence that PD affects interval timing, beat-based rhythm processing, and the temporal organization of voluntary action [26, 40, 41].

This distinction matters because mean signed asynchrony can obscure unstable performance. A participant may tap early on some beats and late on others, producing an average signed asynchrony close to zero despite poor synchronization. In contrast, absolute asynchrony and variability are more sensitive to the consistency of tap-to-beat alignment. The present results therefore support the view that rhythm-based assessments in PD should include measures of precision, stability, and serial structure, rather than relying only on average lead-lag timing [42].

Tempo further shaped the behavioural pattern, which is relevant because cueing rate is a central parameter in rhythm-based interventions. Synchronization was not uniformly better at slower Tempo. Instead, variability of asynchronies was lower at intermediate rates than at fast or slow rates, and timekeeper noise increased as the pacing interval became longer. This finding challenges the simple assumption that slower auditory cues are necessarily easier for people with PD to use and suggests that optimal cueing rates may depend on the balance between external rhythmic support and the individual’s internal timing precision. A likely explanation for this finding is that longer inter-beat intervals require the internal timing process to maintain a temporal representation across a longer delay, allowing variability to accumulate [4, 33, 34, 37]. For PD, this is clinically relevant because cueing rate is a key parameter in rhythm-based interventions, yet the relationship between pacing rate and motor benefit is not straightforward. The present findings suggest that slower pacing is not necessarily easier: longer pacing intervals were associated with higher timekeeper noise, indicating greater demands on internally maintained timing [25].

## Model-based interpretation: phase correction and timekeeper noise

The most informative model-based group differences were reduced phase correction and increased central timekeeper noise. Phase correction captures the degree to which the next response is adjusted after the immediately preceding asynchrony, and can be seen as a measure of the strength of coupling between movements and external events [20]. Lower phase correction in PD therefore suggests that, after a tap occurred too early or too late, the following tap was adjusted less strongly toward the beat. This should be interpreted as a reduction in error-based adjustment, rather than evidence for one specific cause. Several mechanisms could contribute to this pattern, including less precise estimation of the tap-beat asynchrony, reduced weighting of timing errors when planning the next response, weaker coupling between action and the external beat, attentional fluctuations, or differences in response strategy. The present data do not distinguish among these possibilities, but they show that reduced synchronization accuracy in PD was accompanied by weaker trial-to-trial correction of timing errors. This interpretation is consistent with accounts of sensorimotor synchronization as an active error-correction process, in which the timing of each action is continuously adjusted in relation to preceding tap–beat asynchronies [1, 2, 17, 20].

Increased timekeeper noise points to a complementary mechanism. Whereas phase correction concerns how errors are used, timekeeper noise indexes variability in the internal timing process that generates successive response intervals [20]. Higher timekeeper noise in PD suggests that the temporal reference supporting rhythmic action was less precise. This finding fits with the broader view that increased movement variability is a characteristic feature of PD. In classic self-paced finger-tapping work, variability in repetitive movements was increased in PD and could be decomposed into components attributed to central timing and motor implementation processes [43]. The present findings extend this idea to externally paced rhythmic synchronization: participants with PD were not only more variable in their observed taps but showed evidence of reduced precision in the latent timing process from which those taps were generated.

Motor noise, assumed to reflect delays in the peripheral motor system [14], still contributed to synchronization variability in the parameter-prediction models, so execution variability remains relevant. However, the group contrast pointed more strongly to central timekeeping and error correction than to motor-output noise. This pattern is important because clinically visible motor impairment does not reveal which component of timing control is altered. The two-level timing tradition has long emphasized that central timing variability and motor implementation variability can be separated from the serial structure of tapping data [4, 33, 34, 37]. ADAM extends this classical decomposition of movement variability from self-paced tapping to externally paced synchronization. In addition to estimating timekeeper and motor-noise components, the model estimates how strongly each response is adjusted after the preceding tap–beat error. This allowed us to ask whether increased variability in PD reflected noisier motor execution, less stable internal timing, weaker error-based correction, or a combination of these mechanisms. In the present data, the PD pattern was best characterized by higher timekeeper noise and lower phase correction, with no robust global increase in motor noise.

The present task used steady, isochronous sequences, so the adaptation-only version of ADAM was appropriate [20, 27]. This allowed estimation of phase correction, period correction, timekeeper noise, and motor noise under controlled Tempo conditions. It also means, however, that the anticipation and joint-integration components of the full ADAM framework were not directly tested. Future studies using tempo-changing, syncopated, or socially interactive rhythms could test whether the present adaptation findings extend to anticipatory prediction and interpersonal timing mechanisms [22, 27, 31, 44].

## Parkinson’s disease as a model for probing timing in the brain

PD is a progressive movement disorder in which altered timing, action initiation, and gait coordination directly affect everyday motor function. For this reason, PD is not only clinically relevant, but also a useful model for probing the neural architecture of timing. The disorder affects dopaminergic cortico-basal ganglia circuits that are implicated in interval timing, beat-based rhythm perception, action initiation, and the temporal sequencing of movement [26, 40, 41]. Because auditory rhythms can partially support movement in PD, the disorder provides a natural way to study the balance between internally generated timing and externally supported synchronization [12, 45].

The basal ganglia are frequently linked to beat-based timing because they support the extraction and prediction of regular temporal structure [46, 47]. At the same time, timing does not rely exclusively on the basal ganglia, but cerebellar, premotor, supplementary motor, auditory, and parietal systems also contribute to interval timing, temporal prediction, and sensorimotor correction [48–51]. ADAM fits this distributed view because it separates computational functions that may depend on partially distinct neural circuits [27]. The present findings support an account in which PD affects both the precision of an internal timing signal and the online use of recent synchronization errors, consistent with interactions among basal ganglia, cerebellar, and cortical timing networks [24, 52–54].

This model-based approach refines what can be learned from a patient-control comparison. A standard behavioural analysis can show that synchronization differs between groups. A computational model can ask which latent process differs, and whether that process predicts the observed behavioural impairment [20]. In this sense, PD can be used not only to confirm that timing is altered, but also to test candidate theories about how prediction, correction, and variability contribute to rhythmic action.

## Computational markers and clinical perspectives

A central implication of the present work is that ADAM parameters may serve as candidate computational markers of disease-relevant timing mechanisms. We use the term marker cautiously. The present cross-sectional study does not establish diagnostic, prognostic, or treatment-response biomarkers in the formal clinical sense. However, ADAM parameters are objective, quantifiable, and interpretable in relation to candidate mechanisms. Instead of describing a participant as simply better or worse at sensorimotor synchronization, the model estimates whether their performance is characterized by reduced error correction, increased central timing variability, increased motor variability, or altered period updating.

This approach complements the broader movement toward digital and computational phenotyping in PD, where wearable devices, smartphones, and sensor-based motor tasks are increasingly used to obtain objective measures beyond the clinic visit [55–57]. Many digital measures are descriptive, such as tapping speed, step count, gait variability, or tremor amplitude. Model-based rhythmic markers add a mechanistic layer. Two individuals may show similar tapping variability, but one may be dominated by timekeeper noise and another by weak phase correction. Such distinctions could matter if they predict different responses to cueing, medication, training, or neuromodulation.

This is particularly relevant for RAS. RAS is often described as an external temporal scaffold that can improve gait speed, stride length, and cadence in PD [6–8, 39]. Yet clinical effects vary across individuals and protocols. Dalla Bella et al. (2017) [12] provide a direct precedent for linking this variability to phase correction: in their RAS training study, PD participants who showed greater phase correction during an adaptive tapping task were more likely to show a positive gait response to RAS. The present findings extend this logic by showing that phase correction can be estimated from steady-tempo synchronization and is reduced in PD. Although we did not test RAS responsiveness directly, lower phase correction may index a reduced ability to use auditory timing errors to adjust upcoming movements, a mechanism that could be relevant for understanding why some individuals benefit more from rhythmic cueing than others [12]. A rhythmically skilled participant may therefore be one who can use external rhythmic cues not only to follow a beat, but also to correct timing errors efficiently from one movement to the next.

In future work, ADAM-derived parameters could help move rhythm-based rehabilitation toward individualized rhythmic phenotyping. Participants with relatively preserved phase correction may be able to exploit external auditory cues efficiently, while participants with high timekeeper noise may require cueing that reduces reliance on internally maintained timing. Reducing timekeeper noise may be achievable, for example, through denser events (as in music, where sounds can occur between beats), greater beat salience (as in ‘groove’ based music; [13, 58, 59] or training aimed at temporal stability (e.g., using multimodal auditory-visual stimuli and bimanual coordination tasks; [60, 61]).

Tempo selection may also depend on the timing profile: slower cues provide more time for movement execution but require the internal timing signal to be maintained over longer intervals, whereas faster cues provide more frequent external events but may reduce the time available for error correction [1]. These are hypotheses that require direct clinical testing, but they illustrate how computational modelling could guide patient stratification and intervention design.

## Limitations and future directions

Several limitations should be considered. First, the task used steady-tempo pacing sequences. This was appropriate for estimating adaptation-related parameters, but it precluded direct estimation of the full model’s anticipation and joint self-other integration parameters. Tempo-changing, syncopated, metrically complex, or socially interactive rhythms would allow direct tests of whether PD also affects anticipatory prediction and interpersonal timing [62, 63].

Second, finger tapping should not be equated directly with gait . Tapping provides precise millisecond-level measurement and is well suited to computational modelling, but gait also involves bilateral coordination, balance, posture, stride regulation, and interaction with the environment [64–66]. The value of tapping is that it isolates timing mechanisms that may contribute to gait cueing responsiveness. A key next step is therefore to test whether ADAM parameters predict gait outcomes, freezing of gait, RAS responsiveness, or improvements after rhythm-based training.

Third, participants with PD maintained their regular medication regimen, and all were taking levodopa. This increases ecological validity because participants were assessed under conditions closer to everyday functioning, but it limits conclusions about dopamine-dependent mechanisms.

Dopaminergic medication can have complex effects on timing, learning, cue responsiveness, and motor execution [67–69]. Comparing ON and OFF medication states, or relating ADAM parameters to levodopa equivalent daily dose and medication timing, would help clarify whether reduced phase correction and increased timekeeper noise are medication-sensitive.

Fourth, because the present task used steady-tempo sequences and the adaptation-only version of ADAM, it was designed to estimate error correction and timing variability rather than prediction-related parameters. Future studies using tempo-changing or more complex rhythmic sequences could test whether PD also affects the ability to anticipate upcoming changes in rhythmic structure.

Finally, the present analyses were cross-sectional and behavioural [70]. They identify associations between model parameters and synchronization outcomes, but they do not establish causal neural mechanisms or validated clinical biomarkers. Combining ADAM with neuroimaging, EEG, MEG, or deep-brain recordings could test whether phase correction and timekeeper noise map onto distinct neural signatures in auditory-motor, basal ganglia, cerebellar, and premotor networks. Longitudinal and test-retest studies [71] are also needed to determine whether these parameters are stable within individuals, change with disease progression, respond to training, or predict clinically meaningful outcomes.

## Conclusion

In conclusion, participants with PD showed reduced accuracy and stability during synchronized finger tapping with rhythmic auditory pacing sequences. ADAM revealed that these behavioural deficits were associated with lower phase correction and higher central timekeeper noise, indicating impairments in both adaptive error correction and internal timing precision. Timekeeper noise was the clearest predictor of synchronization accuracy, while phase correction and noise parameters jointly accounted for synchronization stability. These findings support a mechanistic interpretation of rhythmic timing impairment in PD and show how model-based analysis can move beyond descriptive performance measures. More broadly, they suggest that computational rhythmic markers may help link basic theories of timing to clinically relevant questions about RAS, individualized cueing, and rhythm-based rehabilitation.

## Statements and Declarations

### Author Contributions

VPN: Conceptualization, Methodology, Software, Validation, Formal Analysis, Investigation, Data Curation, Writing – Original Draft, Writing – Review & Editing, Visualization, Supervision, Project Administration, Funding Acquisition.

VHK: Investigation, Formal Analysis, Writing – Original Draft, Writing – Review & Editing.

KLC: Investigation, Formal Analysis.

JS: Formal Analysis, Writing – Review & Editing.

PV: Conceptualization, Writing – Review & Editing, Funding Acquisition.

PK: Conceptualization, Formal Analysis, Writing – Review & Editing.

### Data availability

The data, materials, and code are available at https://osf.io/szu9v/overview?view_only=4dc41b89cea14bac9656ed1e57279d6f

### Ethics approval

The study was approved by the Institutional Review Board at the DNC (Dansk Neuroforskningscenter), Aarhus University Hospital, Aarhus, Denmark (DNC-IRB-2021-010), and conducted in accordance with the 1964 Declaration of Helsinki and its later amendments.

### Consent to participate

All participants provided written and verbal informed consent before participation.

