## Supplementary Information for "Computational markers of rhythmic timing in Parkinson’s disease"

### Analyses with Tempo interactions

An exploratory fourth set of analyses, the linear mixed effects models included fixed factors of phase correction, period correction, timekeeper noise, and motor noise, their two-way interactions with Group, three-way interactions with Group and Tempo, and participant as a random effects variable. The formula for the linear mixed effect model is as follows:

$$lmer(Asynchrony \sim (Phase\ Correction\ +\ Period\ Correction\ +\ Timekeeper\ Noise\ +\ Motor\ Noise) * Group * Condition\ +\ (1\ |\ Participant))$$

#### a. Signed asynchrony

Relations between mean signed asynchrony and each ADAM parameter in each participant Group and Tempo condition are shown in **Figure S1**. The linear mixed effects model on these data included fixed factors of phase correction, period correction, timekeeper noise, and motor noise, their interactions with Group, Tempo, and Group x Tempo, as well as participant as a random effects variable. Model comparison revealed that this main mixed-effects model fit the data significantly better than a reduced model with only random effects ($\chi 2$ (39) = 225.2, $p < .001$; Log Likelihood = -275.52 (main) vs -388.12 (reduced). In the main model (**Table S1**), only the main effect of Tempo and an interaction between Group and Tempo were statistically significant, consistent with the results reported in section 3.1. However, ADAM parameters did not emerge as significant predictors of signed asynchrony.

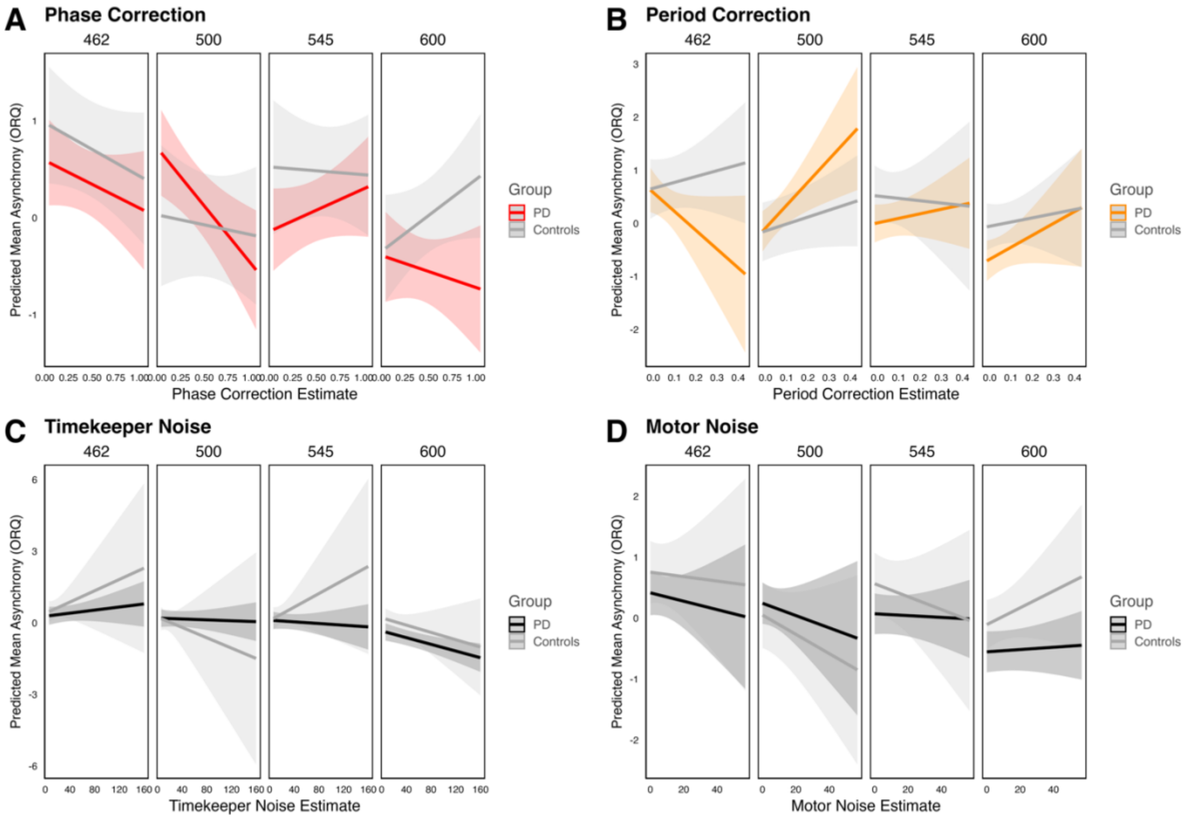


**Figure S1:** Simple slopes of relations between mean signed asynchrony (ORQ-transformed) and each ADAM parameter estimate in each participant Group and Tempo condition. (A) Phase correction; (B) period correction; (C) timekeeper noise; (D) motor noise. PD = Parkinson's Disease.

**Table S1:** Effects of ADAM parameters, Group, and Tempo on mean signed asynchrony. *** p < 0.001; ** p < 0.01; * p < 0.05.

| Model term | Df 1 | Df 2 | *F* value | *p* value |
|---|---|---|---|---|
| Phase correction | 1 | 367.42 | 1.358 | 0.245 |
| Period correction | 1 | 358.37 | 2.050 | 0.153 |
| Timekeeper noise | 1 | 413.91 | 0.001 | 0.975 |
| Motor noise | 1 | 377.93 | 0.662 | 0.416 |
| Group | 1 | 80.98 | 1.059 | 0.306 |
| Tempo | 3 | 354.15 | 21.368 | 0.000*** |
| Phase correction x Group | 1 | 367.42 | 1.072 | 0.301 |
| Phase correction x Tempo | 3 | 357.16 | 2.167 | 0.092 |
| Period correction x Group | 1 | 358.37 | 0.059 | 0.808 |
| Period correction x Tempo | 3 | 356.99 | 2.512 | 0.058 |
| Timekeeper noise x Group | 1 | 413.91 | 0.163 | 0.687 |
| Timekeeper noise x Tempo | 3 | 354.84 | 2.212 | 0.086 |
| Motor noise x Group | 1 | 377.93 | 0.000 | 0.994 |
| Motor noise x Tempo | 3 | 353.98 | 1.403 | 0.242 |
| Group x Tempo | 3 | 354.15 | 3.316 | 0.020* |
| Phase correction x Group x Tempo | 3 | 357.16 | 1.595 | 0.190 |
| Period correction x Group x Tempo | 3 | 356.99 | 2.022 | 0.110 |
| Timekeeper noise x Group x Tempo | 3 | 354.84 | 0.801 | 0.494 |
| Motor noise x Group x Tempo | 3 | 353.98 | 0.446 | 0.720 |

### b. Absolute asynchrony

Relations between mean absolute asynchrony and each ADAM parameter are displayed in **Figure S2**. Model comparison revealed that the main mixed-effects model fit the data significantly better than a reduced random-effects-only model (χ2 (39) = 144.58, $p < .001$; Log Likelihood = -75.42 (main) vs -147.71 (reduced). In the main model (**Table S2**), timekeeper noise estimates predicted absolute asynchrony, with higher timekeeper noise associated with larger mean asynchrony. There was also a significant main effect of Group on asynchrony, and a significant Group x Tempo interaction, corresponding to the effects described in section 3.1 (though in the analyses here, with ADAM parameters, the main effect of Tempo was not statistically significant, perhaps due to its relationship with timekeeper noise). All other effects were non-significant.

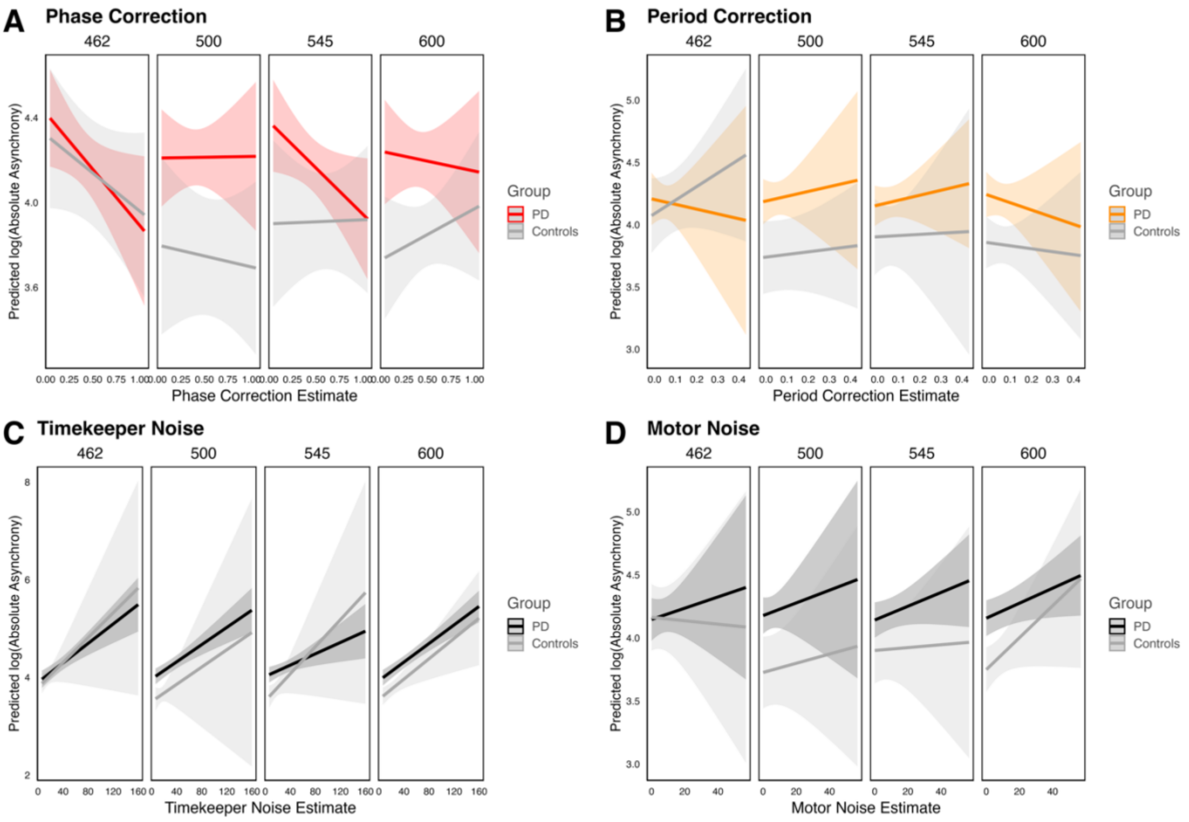


**Figure S2:** Simple slopes of relations between mean absolute asynchrony (log-transformed) and each ADAM parameter estimate in each participant Group and Tempo condition. (A) Phase correction; (B) period correction; (C) timekeeper noise; (D) motor noise. PD = Parkinson's Disease.

**Table S2:** Effects of ADAM parameters, Group, and Tempo on mean absolute asynchrony. *** p < 0.001; ** p < 0.01; * p < 0.05.

| Model term | Df 1 | Df 2 | *F* value | *p* value |
|---|---|---|---|---|
| Phase correction | 1 | 389.43 | 1.967 | 0.162 |
| Period correction | 1 | 369.41 | 0.113 | 0.737 |
| Timekeeper noise | 1 | 412.04 | 15.464 | 0.000*** |
| Motor noise | 1 | 393.50 | 2.197 | 0.139 |
| Group | 1 | 112.43 | 6.702 | 0.011* |
| Tempo | 3 | 357.47 | 1.938 | 0.123 |
| Phase correction x Group | 1 | 389.43 | 0.908 | 0.341 |
| Phase correction x Tempo | 3 | 366.57 | 1.247 | 0.292 |
| Period correction x Group | 1 | 369.41 | 0.211 | 0.646 |
| Period correction x Tempo | 3 | 365.91 | 0.266 | 0.850 |
| Timekeeper noise x Group | 1 | 412.04 | 0.335 | 0.563 |
| Timekeeper noise x Tempo | 3 | 358.51 | 0.065 | 0.978 |
| Motor noise x Group | 1 | 393.50 | 0.037 | 0.847 |
| Motor noise x Tempo | 3 | 358.16 | 0.525 | 0.665 |
| Group x Tempo | 3 | 357.47 | 2.675 | 0.047* |
| Phase correction x Group x Tempo | 3 | 366.57 | 0.374 | 0.772 |
| Period correction x Group x Tempo | 3 | 365.91 | 0.276 | 0.843 |
| Timekeeper noise x Group x Tempo | 3 | 358.5 | 0.233 | 0.873 |
| Motor noise x Group x Tempo | 3 | 358.160 | 0.386 | 0.763 |

### c. Coefficient of Variability

Relations between the CV of asynchronies and each ADAM parameter are shown in Figure 10. Model comparison revealed that the main mixed-effects model fit the data significantly better than a reduced random-effects-only model ($\chi2$ (39) = 639.87, $p < .001$; Log Likelihood = 196.24 (main) vs -123.69 (reduced). In the main model (**Table S3**), phase correction, timekeeper noise, and motor noise emerged as predictors of the CV of asynchronies, each individually and in interactions with Group, as in the analyses reported in section 3.3.1 (which excluded interactions with Tempo). In the current analysis, there was a significant main effect of Tempo, and an interactive effect of Tempo and timekeeper noise on CV of asynchronies. This effect is not straightforward to interpret but might indicate that the relation between timekeeper noise and CV of asynchronies got weaker as Tempo decreased. Although the effect of Group was no longer significant alone, the Group x Tempo interaction was significant, and 3-way interactions with timekeeper noise and motor noise estimates emerged. From **Figure S3**, it appears that the relation between timekeeper noise and CV was more consistent across tempi in PD than controls, whereas the relation between motor noise and CV was more consistent across tempi in controls than PD participants.

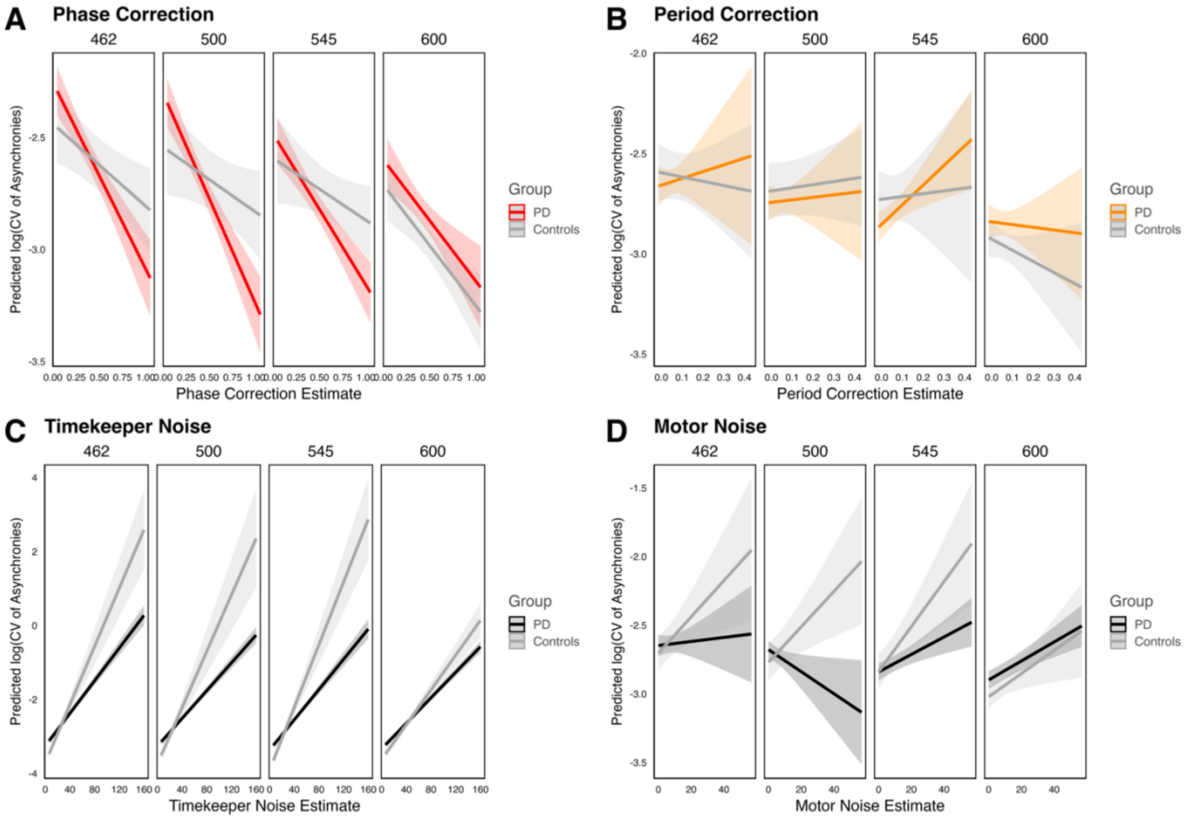


**Figure S3:** Simple slopes of relations between the CV of asynchronies (log-transformed) and each ADAM parameter estimate in each participant Group and Tempo condition. (A) Phase correction; (B) period correction; (C) timekeeper noise; (D) motor noise. PD = Parkinson's Disease.

**Table S3:** Effects of ADAM parameters, Group, and Tempo on the CV of asynchronies. *** p < 0.001; ** p < 0.01; * p < 0.05.

| Model term | Df 1 | Df 2 | *F* value | *p* value |
|---|---|---|---|---|
| Phase correction | 1 | 394.96 | 106.744 | 0.000*** |
| Period correction | 1 | 373.31 | 0.347 | 0.556 |
| Timekeeper noise | 1 | 409.18 | 523.852 | 0.000*** |

| Model term | Df 1 | Df 2 | *F* value | *p* value |
|---|---|---|---|---|
| Motor noise | 1 | 398.39 | 22.919 | 0.000*** |
| Group | 1 | 117.58 | 0.069 | 0.794 |
| Tempo | 3 | 359.37 | 27.266 | 0.000*** |
| Phase correction x Group | 1 | 394.96 | 12.296 | 0.001** |
| Phase correction x Tempo | 3 | 370.06 | 0.449 | 0.718 |
| Period correction x Group | 1 | 373.31 | 1.586 | 0.209 |
| Period correction x Tempo | 3 | 369.22 | 1.119 | 0.341 |
| Timekeeper noise x Group | 1 | 409.18 | 44.737 | 0.000*** |
| Timekeeper noise x Tempo | 3 | 360.52 | 14.187 | 0.000*** |
| Motor noise x Group | 1 | 398.39 | 13.479 | 0.000*** |
| Motor noise x Tempo | 3 | 360.36 | 1.883 | 0.132 |
| Group x Tempo | 3 | 359.37 | 3.969 | 0.008** |
| Phase correction x Group x Tempo | 3 | 370.06 | 1.873 | 0.134 |
| Period correction x Group x Tempo | 3 | 369.22 | 0.326 | 0.806 |
| Timekeeper noise x Group x Tempo | 3 | 360.52 | 6.185 | 0.000*** |
| Motor noise x Group x Tempo | 3 | 360.36 | 2.772 | 0.041* |

**Table S4:** Coefficients for the set of orthogonal planned contrasts testing effects of Group and polynomial trends across Tempo conditions.

| Effect | Contrast | Coefficients | | | | | | | |
|---|---|---|---|---|---|---|---|---|---|
| Group | PD vs Controls | -1 | -1 | -1 | -1 | 1 | 1 | 1 | 1 |
| Tempo | Linear | -3 | -1 | 1 | 3 | -3 | -1 | 1 | 3 |
| | Quadratic | -1 | 1 | 1 | -1 | -1 | 1 | 1 | -1 |
| | Cubic | -1 | 3 | -3 | 1 | -1 | 3 | -3 | 1 |
| Group x Tempo | Group x Linear | 3 | 1 | -1 | -3 | -3 | -1 | 1 | 3 |
| | Group x Quadratic | 1 | -1 | -1 | 1 | -1 | 1 | 1 | -1 |
| | Group x Cubic | 1 | -3 | 3 | -1 | -1 | 3 | -3 | 1 |
